# Single-cell mapper (scMappR): using scRNA-seq to infer cell-type specificities of differentially expressed genes

**DOI:** 10.1101/2020.08.24.265298

**Authors:** Dustin J. Sokolowski, Mariela Faykoo-Martinez, Lauren Erdman, Huayun Hou, Cadia Chan, Helen Zhu, Melissa M. Holmes, Anna Goldenberg, Michael D. Wilson

## Abstract

RNA sequencing (RNA-seq) is widely used to identify differentially expressed genes (DEGs) and reveal biological mechanisms underlying complex biological processes. RNA-seq is often performed on heterogeneous samples and the resulting DEGs do not necessarily indicate the cell types where the differential expression occurred. While single-cell RNA-seq (scRNA-seq) methods solve this problem, technical and cost constraints currently limit its widespread use. Here we present single cell Mapper (scMappR), a method that assigns cell-type specificity scores to DEGs obtained from bulk RNA-seq by integrating cell-type expression data generated by scRNA-seq and existing deconvolution methods. After benchmarking scMappR using RNA-seq data obtained from sorted blood cells, we asked if scMappR could reveal known cell-type specific changes that occur during kidney regeneration. We found that scMappR appropriately assigned DEGs to cell-types involved in kidney regeneration, including a relatively small proportion of immune cells. While scMappR can work with any user supplied scRNA-seq data, we curated scRNA-seq expression matrices for ∼100 human and mouse tissues to facilitate its use with bulk RNA-seq data alone. Overall, scMappR is a user-friendly R package that complements traditional differential expression analysis available at CRAN.

**Highlights:**

- scMappR integrates scRNA-seq and bulk RNA-seq to re-calibrate bulk differentially expressed genes (DEGs).
- scMappR correctly identified immune-cell expressed DEGs from a bulk RNA-seq analysis of mouse kidney regeneration.
- scMappR is deployed as a user-friendly R package available at CRAN.

## Introduction

RNA-seq is a powerful and widely-used technology to measure transcript abundance and structure in biological samples (1). RNA-seq analyses typically compare transcript abundance between conditions by calculating differentially expressed genes (DEGs) (2, 3). When RNA-seq of a whole tissue (bulk RNA-seq) is completed, it is often a challenge to determine the extent to which changes in gene expression are due to changes in cell-type proportion (4). This challenge is addressed by single-cell RNA-seq (scRNA-seq) methods that measure gene expression at a single-cell resolution. Despite many advances, technical limitations (e.g., low gene detection per cell, cell dissociation optimization) and cost currently limit the use of scRNA-seq for hard-to-dissociate cell types and large study designs (5, 6). Importantly, bioinformatics methods that integrate bulk RNA-seq and scRNA-seq demonstrate the highly complementary nature of these two technologies (7–16).

Single cell RNA-seq experiments readily indicate combinations of genes that are involved in the biological functions altered in an experiment or clinical condition. The value of these data is reflected in the growing number of repositories containing publicly available reprocessed scRNA-seq data such as PanglaoDB (17), scRNAseqDB (18), SCPortalen (19), Single Cell Expression Atlas (20) and the Human Cell Atlas (21), which allow for a consistent, tissue-aware reference to the cell-type specificity of individual genes. Indeed, such datasets can be used to interrogate cell-type specific gene expression and enhance bulk RNA-seq analyses in the absence of a matched scRNA-seq experiment (12, 22).

Several methods exist to integrate bulk RNA-seq and scRNA-seq, with the most common class of tools being cell-type deconvolution (12, 14, 15, 23–25). Cell-type deconvolution leverages cell-type specific expression within a scRNA-seq dataset to estimate the relative cell-type proportions within a bulk RNA-seq sample. Estimated cell-type proportions can then be directly compared between conditions to identify alterations in cell-type composition (26, 27). Bioinformatic tools such as csSAM (4) and subsequently released Bseq-sc (28) utilize estimated cell-type proportions to identify DEGs that were not considered differentially expressed with bulk differential analysis alone (2, 3, 29). While powerful, these tools require a larger number of samples than is typically performed in exploratory studies looking for DEGs (4, 28). For this reason, new methods that leverage scRNA-seq to interpret the results from typical bulk-RNA-seq experiments are of value, especially considering the growing number of scRNA-seq reference datasets.

Here we present a tool called single-cell mapper (scMappR) that is designed to infer which cell-types are responsible for DEGs generated using common bulk RNA-seq experimental designs. The purpose of scMappR is to assign cell-type specificity scores to DEGs previously obtained from bulk RNA-seq experiments. Starting with a reference scRNA-seq dataset, scMappR integrates cell-type proportions and cell-type specific expression to compute and visualize the putative cell-type origins of DEGs identified in bulk RNA-seq analysis. We first demonstrate that scMappR can identify validated cell-type specific gene expression by taking advantage of a reference data set (23) where bulk RNA-seq was performed on cell-sorted samples. We show that scMappR can identify bonafide differential gene expression changes emanating from a minority cell population present in the mouse kidney during regeneration (13). Overall, scMappR is a freely available R package (available on CRAN) that provides important cell-type specificity to a set of user-provided DEGs.

## Materials and methods

### R Package: scMappR

We built an R package which we call scMappR to compute and visualize the roles that different cell-types play upon the identification of DEGs. scMappR contains the bioinformatic pipeline to process scRNA-seq data from a count matrix to formats compatible with scMappR. scMappR is currently stored on CRAN (https://cran.r-project.org/web/packages/scMappR/index.html). Reprocessed scRNA-seq cell-type matrices are stored in a separate Github repository (https://github.com/wilsonlabgroup/scMappR_Data).

### Computation and visualization of cell-type contextualized DEGs and cell-type specific pathway analysis

scMappR combines differential expression, cell-type expression, and cell-type proportions to generate cell-weighted fold-changes (cwFold-changes, *cwΔ*). Specifically, scMappR reweighs the fold-changes of bulk DE (*Δ*) genes by the fold-change of cell-type specificity (e.g., cell-type vs. other cell-types) identified in the reference scRNA-seq dataset (*ξ*), and estimated cell-type proportion. These proportions are estimated through RNA-seq deconvolution with the inputted gene’s expression removed from the count and signature matrices. A signature matrix is defined as a gene-by-cell-type matrix populated with the relative expression of a gene in each cell-type. Cell-type proportions (*π*) are estimated with DeconRNAseq (15) and cell-types with >1% of cell-type proportions are used in subsequent analyses (Figure 1). Then, estimated cell-type proportions are made independent from the cell-type expression of each gene using an orthogonalization method based on a leave-one-out approach (30) (Figure 1). Specifically, for each gene, the gene is removed from both the count matrix and the signature matrix (*ξ*). Then, we complete RNA-seq deconvolution with DeconRNA-seq (15) with that gene excluded. This way, the expected cell-type proportions are independent of the cell-type expression of each gene on a per-gene basis. The average cell-type proportion per cell-type (*π*) and the ratio of cell-types proportion between two conditions 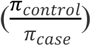 are integrated into scMappR (Figure 1). This reweighting is described in the formula below.

**Figure 1.**
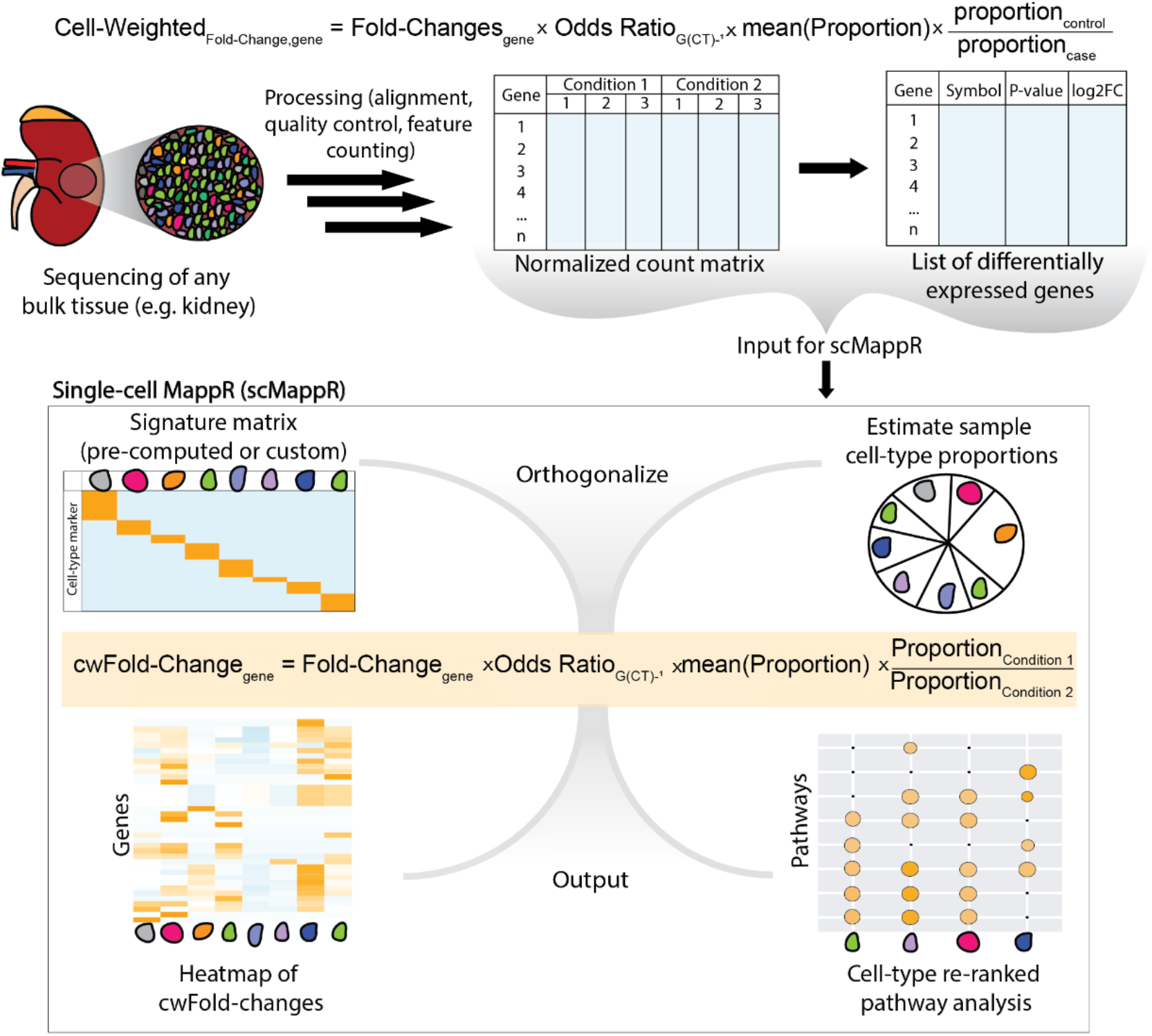
Schematic of the data required to run scMappR and the primary functionalities that scMappR provides. scMappR requires input RNA-seq count data, a list of differentially expressed genes, and a signature matrix (provided by the user or scMappR). For each gene, scMappR then makes cell-type expression independent of estimated cell-type proportions. scMappR then integrates cell-type expression, cell-type proportion, and the ratio of cell-type proportions between biological conditions to generate cell-weighted Fold-changes (cwFold-changes). These cwFold-changes are then visualized (bottom left) and reranked before scMappR computes and plots cell-type specific pathway analyses (bottom right).

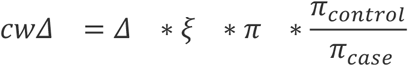

Cell-weighted fold-changes (*cwΔ*) are computed for every DEG in each cell-type. cwFold-changes and endogenous cell-type specificity are then plotted with the Pheatmap R package (31). For every cell-type, each gene on the gene list is reranked by their cwFold-change. Pathway analysis is subsequently completed with g:ProfileR (32) package. By default, scMappR uses the following example command: gProfileR::gprofiler(genes, species, ordered = TRUE, src_filter = c(“GO:BP”, “REAC”, “KEGG”), custom_bg = genes_in_bulk, correction_method = “fdr”) (32, 33) (Figure 1). In this paper we report precision as the g:Profiler summary statistic which g:Profiler defined as the proportion of inputted DEGs that are present in the gene set (32).

### Computation and visualization of the endogenous cell-type specific expression of a gene list

In many instances, it is valuable to gain understanding of the endogenous cell-type expression of a list of genes even when fold-changes and cell-type proportions are not relevant (e.g., significant variants mapping to genes from a Genome Wide Association Study). scMappR plots all of the putative cell-type specific genes in a given signature matrix, as well as the cell-type specific genes that overlap with a user-specified list of human or mouse gene symbols, using the Pheatmap R package (31). scMappR then tests the enrichment of cell-type markers that overlap the user’s list with a Fisher’s exact test (odds ratio > 0, adjusted p-value < 0.05) (34) while using all cell-type markers for every cell-type in that tissue as a background.

### Generation of cell-type signature matrices from publicly available scRNA-seq

Consistently reprocessed scRNA-seq samples were obtained from bulk data in the PanglaoDB (17) project (https://panglaodb.se/samples.html). Briefly, PanglaoDB (17) automatically downloads mouse and human scRNA-seq data before aligning and processing these data in a manner specific to their sequencing platform (Drop-seq, 10X Genomics, and Start-seq) (35, 36). The scMappR package provides the bioinformatic pipeline to convert any scRNA-seq count dataset into a signature matrix with named cell-types within the “process_dgTMatrix_lists” function within scMappR. A signature matrix is defined as a gene-by-cell type matrix containing the likelihood that each gene is expressed in each cell-type. All normalization, clustering, and cell-type maker, and cell-type labelling steps detailed below also describe the “process_dgTMatrix_lists” function and how it was applied to the scRNA-seq data stored in the PanglaDB database (17). To generate signature matrices from scRNA-seq count data, we removed cells with abnormally high mitochondrial content (greater than two standard deviations above the mean in that given sample) (37). Then, normalization, clustering, scaling, and integration of technical replicates were completed using Seurat V3 with the integration anchors feature (38, 39). Cell-type markers are identified using the FindMarkers function in Seurat v3 (default parameters) (38, 39). This function completes differential expression (Wilcoxon’s test as default) between each cell-type and all of the other cell-types. Signature matrices are populated with the rank (-log10(p-value)) to measure enrichment of a cell-type or fold-change output to calculate cwFold-changes. We further use these cell-type markers to define cell-types. Cell-types were identified by extracting the (at maximum) top 30 cell-type markers and converting each gene symbol to human or mouse when necessary using BioMart (40).

Our automated cell-type identification pipeline is based on two gene set enrichment methods, namely the Fisher’s exact test of cell-type markers and Gene Set Variation Analysis (GSVA) of the average expression of each gene per cell-type (34, 41, 42), against two cell-type marker databases, CellMarker and PanglaoDB (16, 17). The CellMarker database manually curated cell-type markers using a literature search of over 100,000 papers and is updated four times per year (16). The PanglaoDB database was generated with a combination of manual curation, co-expression of putative cell-type markers, and community submission (17). scMappR automatically labelled cell-types by appending the most highly enriched cell-type from the CellMarker database to the most highly enriched cell-type using the Panglao database using a two-tailed Fisher’s exact test (16, 17, 34). Cell-types that do not contain significant enrichment with the Fisher’s exact test (34) were labelled unknown, however all cell-types (including unknown) have predicted labels from the GSVA method (41) stored as an output file. Once cell-types were labelled, signature matrices based on rank and fold-change were generated. scMappR reprocesses user-provided scRNA-seq count data with the same pipeline.

We aggregated all the cell-types and cell-type markers into a gene-set database. Each gene-set is designated with the following notation: “SRA ID: tissue: cell-type”. All the cell-type markers within each gene list are consistently processed. This gene-set database can be used for gene-set enrichment using a Fisher’s exact test (34) within scMappR and the gene-set database can be downloaded for other gene-set enrichment analysis tools (33).

The bioinformatic pipeline used to process scRNA-seq from count matrix data is part of the scMappR R package. Users can also provide their own scRNA-seq count matrix, which is converted into a Seurat (39) object that is then processed and converted into a signature matrix using the same methods described above. Users can additionally choose to save intermediary files generated by scMappR to process count matrices into a signature matrix. Specifically, scMappR saves the Seurat object, all cell-type markers, and all possible cell-type labels from both CellMarker and Panglao (using GSVA and the Fisher’s exact test) (16, 17, 34, 38, 39, 41, 42). Finally, the vignette stored in CRAN provides the functions required to convert a Seurat object into a signature matrix. Together, this pipeline can be used as a consistent scRNA-seq processing pipeline from a count matrix of raw scRNA-seq data.

### Processing RNA-seq data from Monaco *et al*., 2019

All fastq files from the peripheral blood mononuclear cells (PBMC) dataset and 29 fluorescence activated cell sorted (FACS) immune cell-types were obtained from GSE107011 (23) using sratoolkit (43). Samples were aligned to the hg38 genome with the STAR aligner (44) using default parameters for paired-end sequencing and filtered for blacklist regions. Reads were assigned to genes using featureCounts (version 1.5.3) with parameters “-s 1 -Q 255 -t exon -O”. Gene models were obtained from GENCODE v33. Counts per million were then calculated for each gene using edgeR and principal component analysis (PCA) was performed (3). Sex differences (N=9 female, 4 male) were measured across the bulk PBMC dataset. Sex differences were also measured in the experiments where RNA-seq was completed after cell-sorting each immune subtype (N = 2 female, 2 male). In both cases, differential expression was completed using DESeq2 (Wald’s test; adjusted P-value < 0.05 and fold-change > 1.5) (2). Cell-type markers were then computed by measuring differential expression of genes in each cell-type against all others (Wald’s test; adjusted P-value < 0.05 and fold-change > 2). Ggplot2 and ggfortify were used to generate all plots (45, 46).

### Processing RNA-seq data from Valle Duraes *et al*., 2020

All fastq files related to RNA-seq on the bulk kidney were downloaded from ArrayExpress (E-MTAB-7957) using wget. These RNAseq bulk kidney samples were aligned to the mm10 genome with the STAR aligner (44) using default parameters for paired-end sequencing and filtered for blacklist regions. Reads were assigned to genes using featureCounts (version 1.5.3) with parameters “-s 1 -Q 255 -t exon -O”. Gene models were obtained from GENCODE M11. Samples were then normalized according to library size and PCA was performed (45, 46). Samples were separated according to strain, sex, condition (i.e. fibrosis or regeneration) and time after injury before differential expression was measured using DESeq2 (Wald’s test; adjusted P-value < 0.05 and fold-change > 1.5) (2).

## Results

### Summary of the scMappR R package and functionality

The scMappR R package contains a suite of bioinformatic tools that provide experimentally relevant cell-type specific information to a list of DEG. The primary function of scMappR is to integrate cell-type expression and cell-type proportions to calculate cell-weighted Fold-changes (cwFold-changes) and cell-type specific pathway analysis from inputted DEGs. cwFold-changes for all genes is ordered within a cell-type to estimate the rank-order of DEGs within each cell-type before a cell-type specific pathway analysis. The cwFold-change for each gene is ordered across cell-types to determine which cell-types were most likely responsible for the differential expression. Investigating cwFold-changes provides context to any differential analysis of a bulk tissue.

scMappR ensures that the cell-type specific expression is relevant to the inputted gene list by containing a bioinformatic pipeline to process scRNA-seq data into a signature matrix, and pre-computed signature matrices of reprocessed scRNA-seq data (Supplementary Table 1) for researchers to choose from. scMappR also provides cell-type specific gene set enrichment of scRNA-seq data for researchers without RNA-seq data and just a gene list.

The function “scMappR_and_pathway_analysis” reranks DEGs to generate cell-type specificity scores called cell-weighted fold-changes (Figure 1, Supplementary Figure 1). Users input a list of DEGs, normalized counts, and a signature matrix into this function. scMappR then re-weights bulk DEGs by cell-type specific expression from the signature matrix, cell-type proportions from RNA-seq deconvolution (15) and the ratio of cell-type proportions between the two conditions to account for changes in cell-type proportion (Figure 1) (See Methods for details).

RNA-seq deconvolution also relies on cell-type specific expression (11, 12, 15, 47). Cell-type specific expression is used to estimate cell-type proportion, making cell-type expression and cell-type proportions dependent values. For each gene, scMappR makes cell-type specific expression and cell-type specific proportion independent values by iteratively removing each gene from the count matrix and the signature matrix before re-calculating cell-type proportions. With cell-type specificity scores computed, scMappR completes pathway analysis by reordering DEGs by their cwFold-changes. Genes that are differentially expressed in the same cell-types move closer in rank, which increases significance in gene set enrichment analysis (32, 41, 48).

With cwFold-changes calculated, scMappR uses two approaches to utilize cwFold-changes to complete cell-type specific pathway analysis. Both approaches are completed by having scMappR rerank DEGs based on their cwFold-changes. Firstly, scMappR reranks DEGs by their cell-weighted fold-change for every cell-type before completing an ordered pathway enrichment. Here, genes are re-ordered by their cell-type specificity scores, but a highly differential ubiquitously expressed DEG may still have a very high rank. Pathway enrichment of the first approach represents biological pathways associated with the rank-change in expression of each cell-type. Secondly, scMappR reranks genes by their increase in cell-type specificity before completing an ordered pathway analysis. For example, if a gene is the 150th most differential DEG in bulk RNAseq and the second most differential cwFold-change for a cell-type, it would have a score of 148 for that cell-type. Pathway enrichment of the second approach represents biological pathways associated with genes most influenced by scMappR.

The “process_dgTMatrix_lists” function in the scMappR package contains an automated scRNA-seq processing pipeline where users input scRNA-seq count data, which is made compatible for scMappR and other R packages that analyze scRNA-seq data (39) (see Methods for details). We leveraged this pipeline to convert over 1,000 scRNA-seq count matrices processed by the PanglaoDB dataset (17) into 245 signature matrices in mouse and human (Supplementary Figure 1).

The functions “tissue_by_celltype_enrichment”, “tissue_scMappR_internal”, and “tissue_scMappR_custom” combine these consistently processed scRNAseq count data with gene-set enrichment tools to allow for cell-type marker enrichment of a generic gene list (e.g. GWAS hits) (see Methods for details). The “tissue_by_celltype_enrichment” function allows for gene set enrichment of all cell-type markers across all cell-types and tissues in scMappR. This gene-set database has the advantage of every cell-type marker originating from a completely consistent bioinformatic analysis. Alternatively, “tissue_scMappR_internal” and “tissue_scMappR_custom” provide a more hypothesis driven approach where researchers can ask if their list of genes are more likely to be expressed in one cell-type compared to other cell-types in the same tissue based on the over-representation of cell-type markers.

### DEG list re-ranked by scMappR reflects cell-type-specific differential expression

scMappR provides the bioinformatic infrastructure to identify which cell-types are likely driving previously identified DEGs from bulk RNAseq. To benchmark scMappR, we chose Monaco *et al*., 2019 (23), a dataset that contained bulk RNA-seq in peripheral blood mononuclear cells (PBMC) (N=13) and RNA-seq of 29 cell-types after cell-sorting (N=4 each) (23). This dataset is ideal to test scMappR as it contains the same biological contrast in bulk RNA-seq data, and in cell-sorted RNA-seq data of the same cell-types. Using full-length RNA-seq of these individual cell-types foregoes the current limitations of differential analysis in scRNA-seq (5, 6) due to issues with dropout and batch effects inherent to scRNA-seq and can thereby be used as an empirical benchmark. This study used males and females in the bulk RNA-seq (N=9 male, 4 female) and in the cell-sorted RNA-seq analyses (N=2 male, 2 female). This allowed for the bulk and cell-type specific measurements of sex differences (Figure 2A, B). We further built a signature matrix using these cell-sorted RNA-seq data by calculating differential expression of each cell-type (sexes combined) vs. all the others.

**Figure 2.**
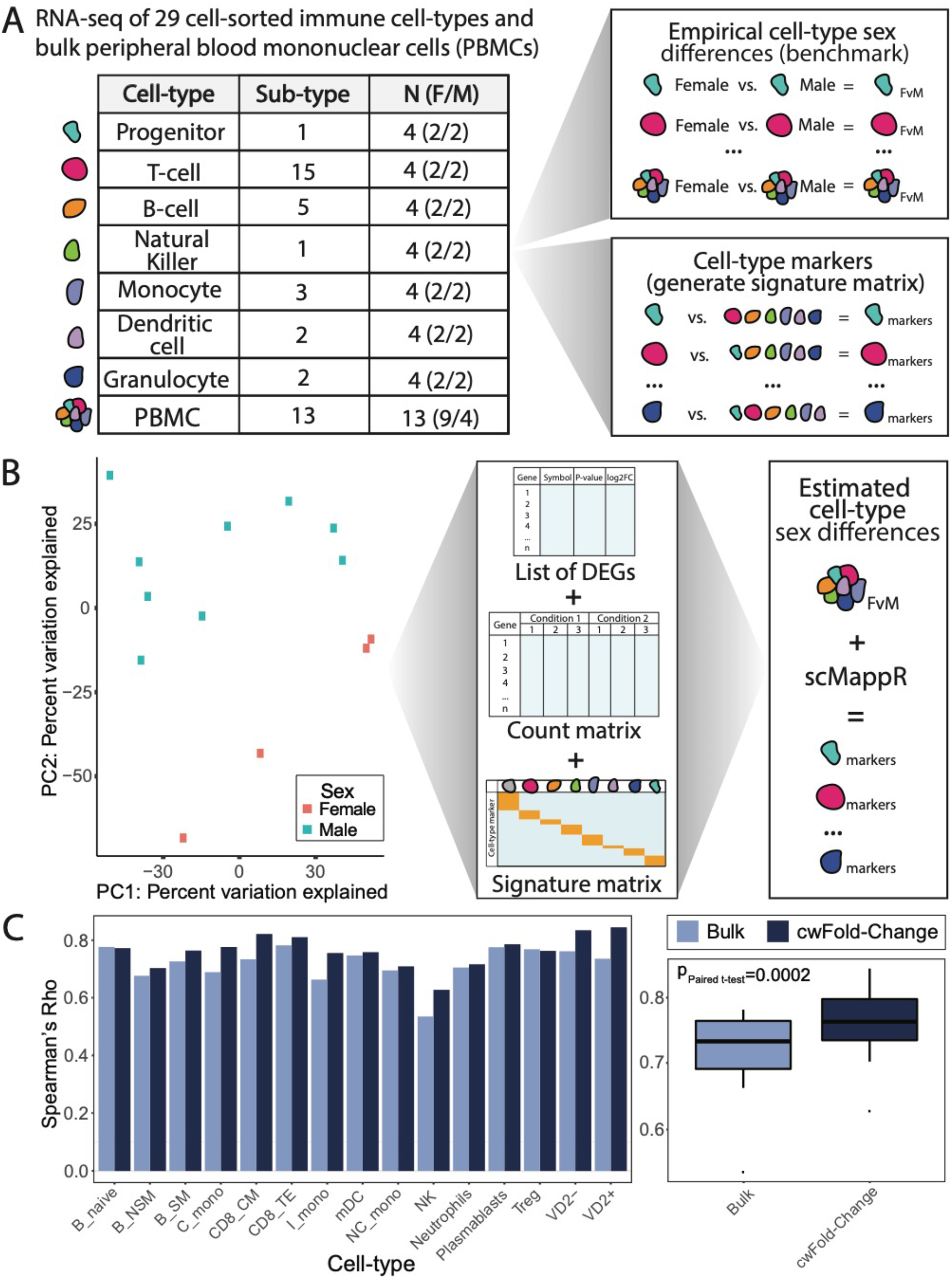
Benchmarking scMappR workflow and results. A) Overview of samples and cell-types from Monaco et al., 2019. Sex differences within each cell-type are computed and the cell-type specific fold-changes in the genes that are differentially expressed in the peripheral blood mononuclear cells (PBMC) dataset are used. Each column of the signature matrix is the fold-change of expression from each cell-type against all of the other cell-types and each row is a cell-type marker. B) Overview of how scMappR was used to estimate cell-type specific sex differences from PBMCs. Principal component analysis shows linear separation of male and female PBMC samples. Differentially expressed genes derived from computing sex differences, the normalized count matrix, and signature matrix generated in (A) were inputted into scMappR. C) Improvement that cell-weighted fold-changes (cwFold-changes) have on cell-type specificity for every cell-type measured with a bar chart. Dark bars are the correlation cwFold-changes with cell-type specific fold-changes. Light = bars are the correlation between cell-types (left) and a boxplot of the correlations across cell-types (right). Improvement in correlation is measured with a one-tailed paired Student’s t-test. *Bulk/PBMC = Peripheral Blood Mononuclear Cells, Neutrophils = Neutrophils, Progenitor = Progenitor, Basophils = Basophils, pDC = Plasmacytoid dendritic cells, Plasmablast = Plasmablast, mDC = myeloid dendritic cells, B_naive = naive B cells, NC_mono = non-classical monocytes, C_mono = classical monocytes, MAIT = MAIT cells, B_SM = Switched memory B cells, VD2- = non-Vd2 gd T-cells*.

Through bulk RNA-seq analysis, we identified 59 DEGs between sexes in PBMCs (Wald’s test; adjusted P-value < 0.05 and fold-change > 1.5) (Supplementary Figure 1A). RNA-seq deconvolution tools including DeconRNA-seq function optimally when there are more samples than cell-types (15, 47). Therefore, we tested scMappR with the top 12 most variable cell-types (one fewer than the 13 bulk samples). We then tested if the fold-changes of these 59 sex-biased DEGs were more highly correlated to the same 59 genes in these 12 cell-types using Spearman’s correlation (Figure 2C). We found that for every cell-type, scMappR’s cwFold-changes either increased or made no change to cell-type specificity (average rho increase = 0.0471, one-tailed Paired Student’s t-test, p = 2.00 x 10^−4^). Overall, scMappR significantly increased the cell-type specificity of a study that already contained a high correlation between cell-type specific DEGS and bulk cell-type specific differential expression (rho = 0.535-0.777). The high correlation between bulk DEGs and cell-type specific DEGs are explained in part by 16 of these genes mapping to the Y chromosome (Supplementary Figure 1B). We then removed the Y chromosome genes due to their inherent sex-biased gene expression and repeated this analysis. We found the same improvement of cell-type specificity (average rho increase = 0.0660, one-tailed Paired Student’s t-test, p = 0.0153), showing that ubiquitously expressed DEGs do not improperly influence scMappR’s cwFold-changes. Removing these DEGs did decrease the correlation between bulk DEGs and cell-type specific DEGs (0.708 with Y chromosome genes to 0.430 without). Together, we show that genes that are ubiquitously differentially expressed do not influence scMappR but do influence the baseline correlation between bulk DEGs and empirically measured cell-type specific DEGs.

We next tested whether scMappR is robust to any combination of cell-types, and not just the 12 most variable cell-types. Monaco *et al*., 2019 contained 13 bulk RNA-seq (PBMC) samples, allowing us to test scMappR using 12 cell-types at once (15, 47). We randomly sampled 12/29 cell-types and re-calculated the p-value and change in correlation for 100 permutations to ensure that our results are not biased by the cell-types that we selected. This permutation-based analysis showed that regardless of the cell-types selected, there was always a statistically significant increase in cell-type specificity (mean P-value = 1.83 x 10^−4^, mean Rho increase = 0.0545) (Supplementary Figure 2). scMappR’s cwFold-changes improved cell-type specificity of individual genes in two ways. Firstly, scMappR increased the rank of differentially expressed cell-type markers (Supplementary Figure 3). Secondly, scMappR decreased the rank of DEGs that were not expressed in a particular cell-type (Supplementary Figure 3). Together, this analysis showed that scMappR can significantly improve the correlation of bulk DEGs to cell-type specific DEGs.

### scMappR reveals cell-type specific DEGs during mouse kidney regeneration

After benchmarking scMappR, we tested how scMappR can be used to assign cell-types contributing to DEGs generated from a representative, well-designed bulk RNA-seq study of a heterogeneous tissue. To do this, we reanalyzed data from Valle Duraes *et al*., 2020 (13) who interrogated gene expression changes involved in mouse kidney regeneration before and after injury (13). Kidney regeneration involves multiple cell-type specific processes (49–52), and importantly Valle Duraes *et al*., 2020 used bulk RNA-seq in conjunction with histopathology, cell sorting, and scRNA-seq to implicate T-Cell recruitment as a critical part of the regeneration process (13). We reasoned that this is an ideal model RNA-seq study to showcase scMappR as Valle Duraes *et al*., 2020 is well-powered, and includes detailed experimental follow-up of cell-type specific responses (13).

Their bulk RNA-seq study design (13) has 50 total samples split into fibrosis (using wild-type mice) and regeneration (using B6.Cg-Foxp3tm2(EGFP)Tch/J mice) models after injury (days 0, 3, 7, 14, 28 and 42) (N=3-4 per condition/timepoint) (Supplementary Figure 4). For simplicity, we focused on the comparison of the initial two timepoints as these contained the most dramatic changes (day 0 (‘naive’) vs day 3 (injury induced ‘regeneration’)). For every comparison, all samples were used in the RNA-seq deconvolution step of scMappR’s generation of cwFold-changes (all time periods in regeneration and fibrosis). In conjunction, a kidney scRNA-seq dataset from *Tabula Muris*, 2018 (53) was preprocessed and stored in scMappR. We then used scMappR to identify which cell-types are involved in kidney regeneration using both bulk and scRNA-seq datasets.

After reprocessing data in Valle Duraes *et al*., 2020 (13), we identified 2855 DEGs between the ‘naive’ and ‘regeneration’ groups. We found that 394 of these DEGs were kidney cell-type markers in *Tabula Muris*, 2018 (53) (Figure 3A). Using scMappR, we then asked which cell-types had the highest cwFold-changes in DEG comparisons between naïve day 0 and regeneration day 3 groups in the whole kidney. We found clear signatures of fibroblasts, smooth muscle, and endothelial cells, all of which have well-documented roles in kidney regeneration (49–52) (Figure 3B). A subset of immune (“Macrophage, dendritic”) specific DEGs were also found (Figure 3B, Table 1). The immune-specific DEGs were less prevalent than other cell types (Figure 3B), likely due to a lower proportion of immune cells in the kidney (54).

**Table 1.** Over-representation of cell-type markers of consistently processed scRNA-seq data in over 100 mouse tissues when inputting 34 T-cell markers.

| SRA ID | Tissue | Label from<br>CellMarker database | Label from<br>Panglao database | Number of cell-type<br>markers | Number of overlapping<br>cell-type markers | Odds ratio | Adjusted<br>p-value |
| --- | --- | --- | --- | --- | --- | --- | --- |
| SRA653146 | Trachea | Lymphocyte | Nuocytes | 122 | 15 | 83.1 | $2.47 \times 10^{-18}$ |
| SRA667466 | Cortex 3 | Lymphocyte | Nuocytes | 122 | 15 | 83.1 | $2.47 \times 10^{-18}$ |
| SRA653146 | Muscle | Myeloid cell | Natural killer cells | 88 | 13 | 90.6 | $1.08 \times 10^{-16}$ |
| SRA667466 | Dorsal midbrain | Myeloid cell | Natural killer cells | 88 | 13 | 90.6 | $1.08 \times 10^{-16}$ |
| SRA748166 | Cardiac tissue | T cell | Natural killer cells | 132 | 13 | 60.2 | $9.58 \times 10^{-15}$ |
| SRA801845 | Cardiac progenitor<br>cells | T cell | Natural killer cells | 132 | 13 | 60.2 | $9.58 \times 10^{-15}$ |
| SRA638923 | Small intestine | Immune cell | Natural killer cells | 92 | 11 | 66.9 | $4.40 \times 10^{-13}$ |
| SRA711739 | Embryonic<br>fibroblasts | Epithelial cell | Natural killer T cells | 60 | 10 | 89.5 | $4.40 \times 10^{-13}$ |
| SRA757237 | Bone marrow | Epithelial cell | Natural killer T cells | 60 | 10 | 89.5 | $4.40 \times 10^{-13}$ |
| SRA653146 | Spleen | T cell | Thymocytes | 68 | 10 | 78.9 | $1.03 \times 10^{-12}$ |

**Figure 3.**
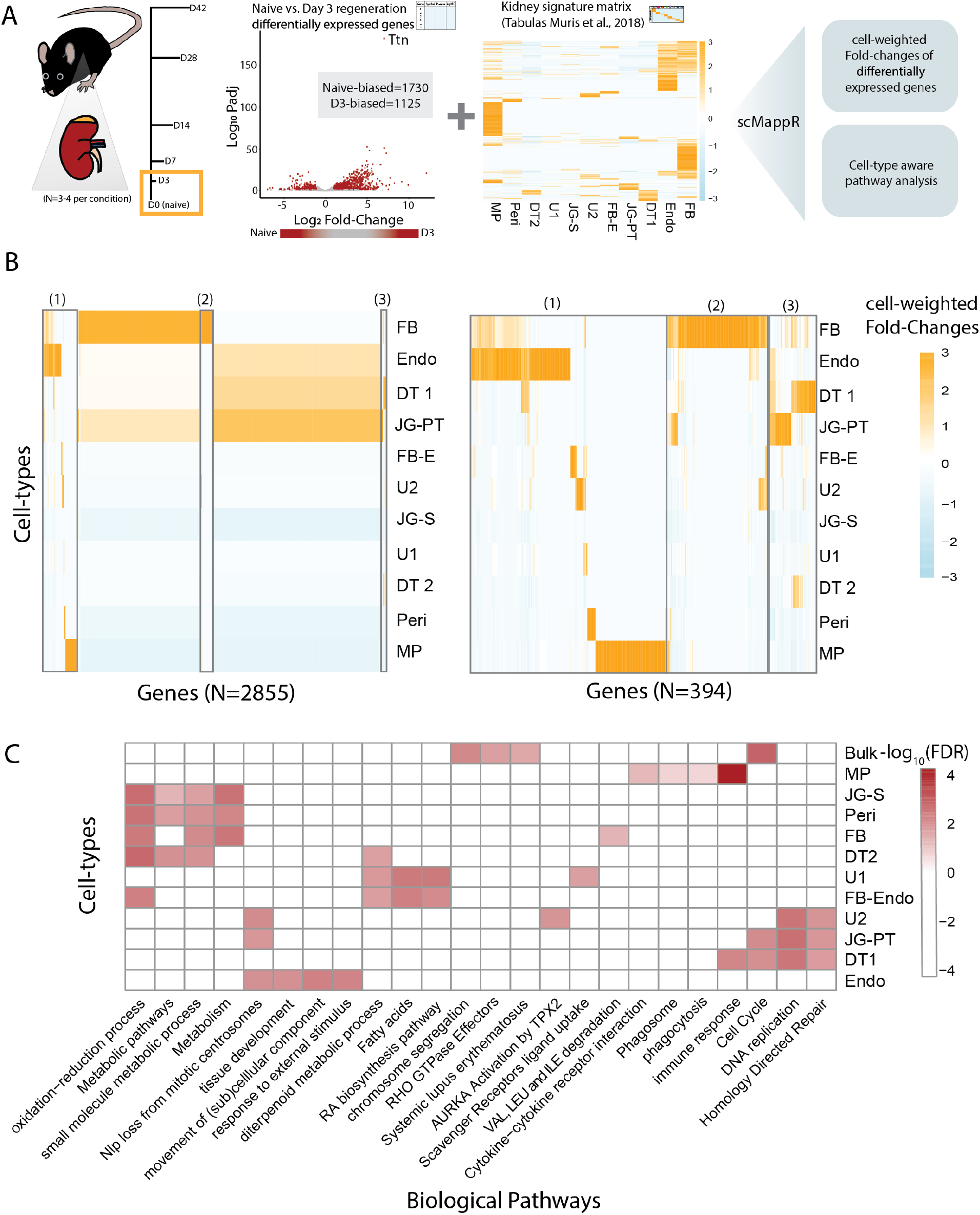
Application of scMappR to identify which cell-types are responsible for differentially expressed genes in kidney regeneration. A) Valle Duraes *et al*., 2020 completed RNA-seq of C57BL/6J mice kidneys at naive (day 0) and multiple timepoints of kidney regeneration post-injury. Between naïve and regeneration day 3 comparisons (shown here), we identified 2855 significantly differentially expressed genes. We then used scMappR to compute cwFold-changes. The normalized count data, the list of differentially expressed genes, and a signature matrix were inputs for this analysis. We used normalized count data from all samples, differentially expressed genes from naïve vs kidney regeneration (naïve (day 0) vs day 3 comparison shown here), and the signature matrix from scRNA-seq in the kidney completed by *Tabula Muris, 2018*. B) Heatmap of gene normalized cwFold-changes of all 2855 differentially expressed genes (left) and the 394 differentially expressed genes that are also identified as cell-type markers in *Tabula Muris*, 2018 (right). The heatmaps on the left and right were produced in the same way except that in the heatmap on the right the genes are filtered for cell-type markers in *Tabula Muris*, 2018. C) A cell-type normalized matrix of the top four most enriched pathways from cell-type specific pathway analysis. For each cell-type, genes were reranked by their increase in cell-type specificity before pathway analysis was completed. *Bulk = bulk kidney, MP = Macrophage, Dendritic, JG-S = Juxtaglomerular, Stem, Peri = Pericyte, FB = Fibroblast, DT2 = Distal Tubule 2, U1 = Unknown 1, FB-Endo = Fibroblast-Endothelial, DT1 = Distal Tubule 1, JG-PT = Juxtaglomerular, Proximal tubule, U2 = Unknown 2, Endo = Endothelial*.

All cell-type labels were identified using the automated cell-type naming process in scMappR and the immune cluster was automatically given the cell-type label “Macrophage, dendritic”. This cell-type contains 430 cell-type markers that enrich for many immune related processes (immune system processes: precision = 0.537, one-tailed hypergeometric test adjusted p-value = 3.65 x 10^−87^; innate immune response: precision = 0.223, one-tailed hypergeometric test adjusted p-value = 1.78 x 10^−38^; adaptive immune response: precision = 0.184, one-tailed hypergeometric test adjusted p-value = 7.95 x 10^−36^; T-cell activation: precision = 0.161, one-tailed hypergeometric test adjusted p-value = 2.10 x 10^−32^). Furthermore, the original *Tabula Muris, 2018* study labelled this cell-type population as “Macrophage and Natural Killer” (53). Interestingly, many cells within this population contain a high expression of naive T-cell markers like *Ccr7* and *Nkg7* (13, 53). These results are unsurprising as T-cells are present in the uninjured kidney (55). Therefore, although this cluster was given the “Macrophage, dendritic” label, it might be better interpreted as a cell-type representing the heterogeneous immune-cell population in the *Tabula Muris*, 2018 (53) kidney.

Overall, the top five most significant pathways of these reranked DEGs showed a common regeneration phenotype across different cell-types at the pathway level (Supplementary Figure 5). For each cell-type, between 52-59% of the pathways were shared between the enriched pathways derived from bulk differential expression compared to pathways derived from genes reranked by cwFold-changes (Supplementary Table 2). Pathways that were only identified in the cell-type specific pathway analyses but not in bulk pathway enrichment were biologically relevant. One such pathway is the “Immune System” gene ontology, which was not significantly enriched with the bulk DEG list but was highly enriched when reranking the same DEGs but by their “Macrophage, dendritic” cwFold-changes (adjusted p-value = 1.62 x 10^−6^). The top five most significant pathways identified by ordering genes based on their rank-change between bulk DEGs and cwFold-changes (Supplementary Figure 6) were related to their cell-type, including significant enrichment of immune related pathways in the “Macrophage, dendritic” cell-type (immune response: precision = 0.125, intersection of DEGs and pathway = 156 genes, one-tailed hypergeometric test adjusted p-value = 6.67 x 10^−19^, cytokine-cytokine receptor interaction: precision = 0.0320, intersection of DEGs and pathway = 37 genes, one-tailed hypergeometric test adjusted p-value = 3.01 x 10^−7^, phagosome: precision = 0.0240,intersection of DEGs and pathway = 30 genes one-tailed hypergeometric test adjusted p-value = 1.82 x 10^−5^, and phagocytosis: precision = 0.126 intersection of DEGs and pathway = 30 genes, one-tailed hypergeometric test adjusted p-value = 3.32 x 10^−5^) (Figure 3C). Taken together, scMappR increases the rank of cell-type specific DEGs, thus allowing for biologically relevant cell-type specific pathway analysis.

In our bulk RNA-seq analysis, we identified three genes, *Il1rl1, Rgs16*, and *Ccr7* as DEGs when naive day 0 and regeneration day 3 were compared. Valle Duraes *et al*., 2020 used three genes as T-cell markers in the CD4^+^ sorted scRNA-seq experiment of naive, regenerating, and damaged kidney (13, 56). We identified an increase in the rank-order of these three DEGs between bulk RNA-seq and “Macrophage, dendritic” cwFold-changes (P-value one-tailed Wilcoxon’s test = 0.047; *Il1rl1: bulk rank 1545; “Macrophage, dendritic” cell rank 1029; Rgs16: bulk rank 354, “Macrophage, dendritic”* = 235; *Ccr7: bulk rank* = 1926, *“Macrophage, dendritic”* rank 367). Notably, when scMappR was applied to the RNA-seq of bulk kidneys and the scRNA-seq of the entire kidney, we were still able to uncover a cell-type specific role of DEGs in a cell-type present in <5% of the bull kidney population (Supplementary Figure 2).

To complete the deconvolution step of scMappR’s generation of cwFold-changes described above, we used 50 total samples across all experimental timepoints (days after injury: 0, 3, 7, 12, 28, 42). However, many studies do not have this number of samples. To investigate whether we would reach similar conclusions using a sample number more representative of routine RNA-seq studies, we repeated our analysis exclusively with the ‘naive’ day 0 (N=4) samples and ‘regeneration’ day 3 (N=3) samples, resulting in seven total biological samples instead of 50. Since scMappR’s generation of cwFold-changes relies on RNA-seq deconvolution, which assumes that there are more samples than cell-types (15, 47), we chose to analyze four cell-types, “proximal tubule, juxtaglomerular”, “endothelial”, “macrophage, dendritic”, and “fibroblast” as they showed variable cwFold-changes at the pathway level (Figure 3, Supplementary Figure 5).

We generated cwFold-changes between ‘naive’ day 0 (N=4) samples and ‘regeneration’ day 3 (N=3) conditions on “proximal tubule, juxtaglomerular”, “endothelial”, “macrophage, dendritic”, and “fibroblast” cell-types using all 50 samples and again with seven samples. When we compared the cwFold-changes between the 50 and 7 sample analyses we found that the rank of cell-type specificity score did not change for any of the cell types (Supplementary Table 3) and that across cell-types, all DEGs maintained the same rank-order of cwFold-changes. The average cell-type proportions for each cell-type was not significantly different between the 50 sample and 7 sample datasets (Supplementary Table 3). Thus, we used a study design with many biological samples to show that scMappR functions appropriately with a sample number representative of routine RNA-seq studies.

### scMappR: projection of a generic gene list onto scRNA-seq data

In addition to disentangling the cell-type specific role of bulk DEGs, scMappR can facilitate the understanding of cell-type specific expression in any list of genes. We tested the cell-type enrichment for the 2855 DEGs measured between naïve day 0 and regeneration day 3 in the kidney across all of the cell-types and cell-type markers stored in scMappR (See Methods). The top ten most significantly enriched cell-types were “proliferating cells and gamma delta T cells” (Supplementary Table 4). To characterize the CD4^+^ scRNA-seq dataset, Valle Duraes *et al*., 2020 (13) utilized a curated set of 34 T-cell marker genes (56). We asked if scMappR in combination with our uniformly processed scRNA-seq data would also consider these as T-cell marker genes. Reassuringly, of the top ten most enriched cell-types, all ten were immune cell-types and four out of ten were from cell-types labelled as T-cells (Table 1).

In addition to testing lists of genes across compendiums of scRNA-seq data, scMappR is useful for interrogating a specific, biologically relevant tissue. This approach is valuable when users have a list of genes from a particular tissue but cell-type proportions cannot be integrated with scRNA-seq expression (e.g. genes mapping to ChIP-seq peaks) (57, 58). As an example, we compared the 2855 DEGs between naive kidney and kidney regeneration (3 hours post injury) against the *Tabula Muris*, 2018 *(53)* kidney scRNA-seq study. We found an over-representation of the immune (“Macrophage Dendritic”) cell-type in the upregulated (regeneration biased) DEGs (FDR adjusted p-value = 1.43 x 10^−5^, odds-ratio = 1.86) and an underrepresentation of the same cell-type in the downregulated (naive baised) DEGs (FDR adjusted p-value = 4.23 x 10^−5^, odds-ratio = 0.33) (Supplementary Table 5). Since the 34 T-cell markers exclusively enriched for the immune (“Macrophage, dendritic”) cell-type (FDR adjusted p-value = 0.00115, odds-ratio = 20.9) (Table 2), we suggest that scMappR did detect evidence of T-cell infiltration which Valle Duraes *et al*., 2020 experimentally validated in their study.

**Table 2.** Over- and under-representation of kidney cell-type markers from scRNA-seq data generated by *Tabula Muris*, 2018 when inputting 34 T-cell markers.

| Cell Type | Total number of cell-type markers in *Tabula Muris*, 2018 | Odds ratio | Adjusted p-value | Number of genes | T-cell marker genes |
| --- | --- | --- | --- | --- | --- |
| Macrophage, dendritic | 430 | 20.9 | 0.00115 | 10 | Klf2,Rgs2,Ccl4,Cd83,Nkg7,Ccl5,Ccr7,Sell,Ifng,Cd7 |
| Endothelia | 560 | 0.138 | 0.169 | 1 | Klf2 |
| Fibroblasts | 548 | 0.317 | 0.793 | 2 | Klf2,Gata3 |
| Distal tubule | 48 | 2.67 | 0.927 | 1 | Gata3 |
| Proximal tubule, juxtaglomerular | 23 | 0.00 | 1.00 | 0 |  |
| Distal tubule1 | 111 | 0.00 | 1.00 | 0 |  |
| Unknown | 49 | 0.00 | 1.00 | 0 |  |
| Pericyte | 49 | 0.00 | 1.00 | 0 |  |
| Fibroblast, endothelia | 43 | 0.00 | 1.00 | 0 |  |
| Unknown1 | 3 | 0.00 | 1.00 | 0 |  |
| Stem, juxtaglomerular | 23 | 0.00 | 1.00 | 0 |  |

Overall, our results show that scMappR can calibrate genes from a representative RNA-seq study design and detect biologically relevant cell-type specific enrichments from gene lists using compendiums of scRNA-seq data. To facilitate further testing on addition RNA-seq data sets and gene lists we made scMappR freely available as an R package on CRAN.

## Discussion

scMappR is an R package designed for the primary purpose of estimating which cell-types contribute to a list of DEGs from bulk RNA-seq. scMappR integrates both cell-type expression and cell-type proportion to generate cell-type specificity scores (cwFold-changes). scMappR’s cwFold-changes applied to bulk DEGs were correlated to empirically measured DEGs compared to bulk DEGs alone (Figure 2). Computing cwFold-changes on DEGs across kidney regeneration allowed for the measurement of which cell-types are responsible for which DEGs and for cell-type specific pathway analysis (Figure 3). scMappR should provide valuable cell-type specificity to a list of DEGs, and scMappR can be performed in many experimental contexts.

The general usability of scMappR with bulk RNA-seq analysis is facilitated in two ways. Firstly, scMappR stores consistently processed mouse and human signature matrices for researchers to choose from. Secondly, scMappR contains the bioinformatic pipelines that allow researchers to reprocess any scRNA-seq count dataset into a signature matrix. There are thousands of viable scRNA-seq processing pipelines (59), and to accommodate this, scMappR allows researchers to input their own signature matrix, scRNA-seq count data, or processed scRNA-seq dataset. From there, scMappR has functions to convert this date into a signature matrix compatible with scMappR’s cwFold-change generation.

We tested the functionality of scMappR by comparing bulk DEGs to empirically measured cell-type specific DEGs before and after scMappR was applied. scMappR’s cwFold-changes were more correlated to cell-type specific DEGs than bulk DEGs (Figure 2). This benchmark suggests that scMappR’s cwFold-changes increase cell-type specificity and that our assumption that the cell-type of origin of a DEG is derived from the cell-type where the gene is most highly expressed is a reasonable assumption. However, it is important to note that scMappR is limited by the initial DEG list itself. For example, a gene that is upregulated in one cell-type and downregulated in another will not be differentially expressed and hence not be part of a scMappR analysis. Unlike scMappR, methods such as BSeq-sc (28) use estimated cell-type proportions as a covariate of differential analysis before applying csSAM, a least-squares regression and empirical FDR (4), to discover DEGs that were undetectable by traditional bulk RNA-seq differential analysis. Importantly, BSeq-sc, requires many biological samples (i.e. 82 samples to discover novel DEGs in three cell-types (28)) which makes these approaches complementary to scMappR depending on the researchers dataset.

scMappR relies on RNA-seq deconvolution to generate cwFold-changes and therefore follows the same assumptions and limitations of RNA-seq deconvolution. Important RNA-seq deconvolution assumptions related to scMappR’s cwFold-changes are that there should be more samples than cell-types (47) and that RNA-seq deconvolution assumes that that the cell-types within a signature matrix make up the entire bulk sample. Limitations of RNA-seq deconvolution related to scMappR are that RNA-seq deconvolution is sensitive to the number of samples, the number of cell-type markers, the processing of the bulk RNA-seq and normalization of the scRNA-seq data (60). While in principle any deconvolution method that relies on a signature matrix would work, we chose to use DeconRNA-seq (15) in the RNA-seq deconvolution step of scMappR’s cwFold-change generation because of its computational efficiency and the size of the input signature matrix. DeconRNA-seq allows for signature matrices upwards of 3,000 genes, and can identify cell-type proportions of ten cell-types in 50 samples in less than three seconds (15). This computational efficiency is important to scMappR because, for example, if there are 2,500 DEGs, the cell-type deconvolution is completed 2,501 times. Furthermore, scMappR’s cwFold-changes rely on the average relative cell-type proportions across conditions and thus it is reasonable to sacrifice a small amount of sensitivity in cell-type proportion estimation for computational efficiency (11, 12, 14, 47).

scMappR leverages scRNA-seq data to characterize the cell-type specificity of a list of bulk DEGs while providing a cell-type marker database to test the over-representation of cell-type markers in any gene list. Currently, single-cell genomic technologies are evolving and expanding to include new assays such as single cell open chromatin (single cell ATAC-seq) (61, 62) and single cell DNA methylation (DNAm) (63, 64), scRNA-seq across many biological conditions with replicates, and single-cell genomics with fewer technical limitations. As these methodologies improve, tools like scMappR that aid in integrating bulk and single-cell differential genomics will become increasingly important.

In summary, we have shown that scMappR can accurately estimate which cell-types contain DEGs. scMappR also has the potential to uncover biological signals that may have otherwise been masked in traditional bulk differential analysis. The scMappR method is stored in a user-friendly R package that provides supplementary pipelines to support researchers with diverse experimental designs and sample sizes. Overall, scMappR should be easy to incorporate into existing RNA-seq pipelines and serve as a facile way to incorporate scRNA-seq data into differential gene expression analyses.

## Data availability

The scMappR R package is available at CRAN (stable release) https://cran.r-project.org/web/packages/scMappR/index.html. The scMappR developmental version is available on github https://github.com/wilsonlabgroup/scMappR_Data. All code and files to generate figures and tables can be found on figShare (preprint link) https://figshare.com/s/3b5cfb597a0b3bc2801c.

## Supplementary Data

Supplementary figures and tables are available at NAR online.

## Author’s Contributions

D.S., M.F.M., A.G., and M.W. conceived of the method described in this manuscript. D.S. created the R code and the R package with support from H.Z. Troubleshooting and methodological utility and testing of scMappR tool was completed by D.S., M.F.M., L.E., H.H. The manuscript was written by D.S., M.F.M., C.C., A.G., and M.W. with support from all authors. M.H., A.G. and M.W supervised the work.

