## Supplementary_Material for "Single-cell mapper (scMappR): using scRNA-seq to infer cell-type specificities of differentially expressed genes"

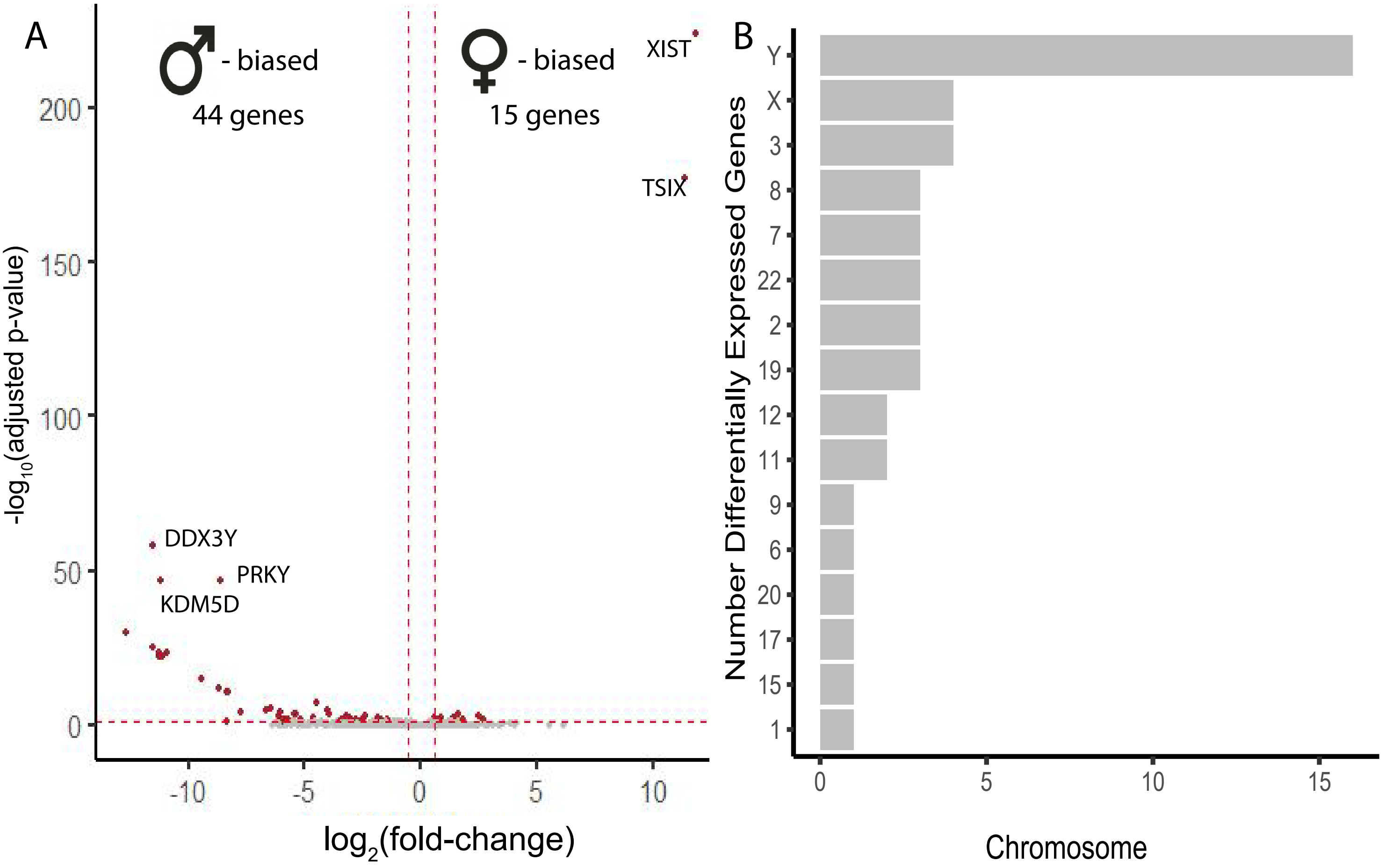


**Supplementary Figure 1. Summary of sex biased differentially expressed genes in Peripheral Blood Mononuclear Cell (PBMC) samples sequenced in Monaco et al., 2019.** A) Volcano plot of the 59 sex differences. Each point is a different gene. The X-axis shows the log2(fold change) of each gene, the Y-axis shows the -log10(Padj) of each gene. Genes in grey are not differentially expressed and genes in red are differentially expressed (Wald's test; Padj < 0.05 and fold change > 1.5). B) Barplot of different chromosomes to where sex differences map on the human genome.

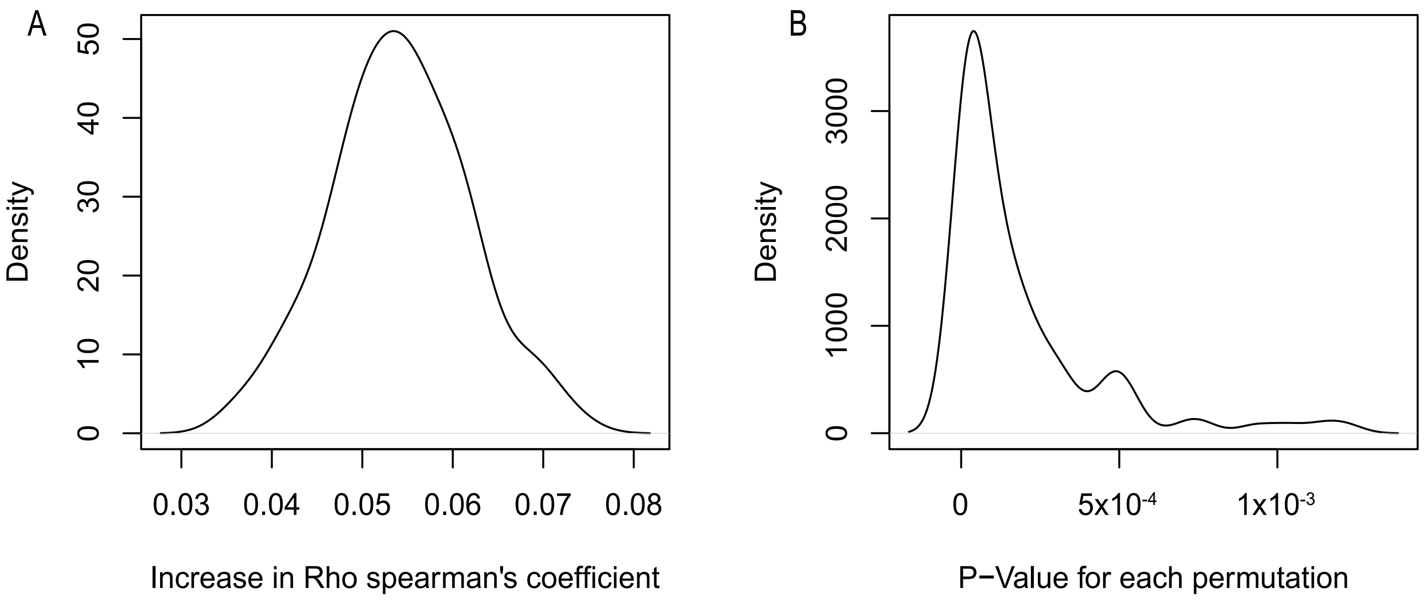


**Supplementary Figure 2. Density plot of the average Spearman's rho increase and the paired Student's T-test P-value when permuting 12 random cell-types.** A) The X-axis is the average increase in Spearman's Rho before and after scMappR is applied and the Y-axis is the density of the increase in Rho after each permutation. B) The X-axis is the average P-value from a paired Student's T-test and the Y axis is the density of the P-value after each permutation.

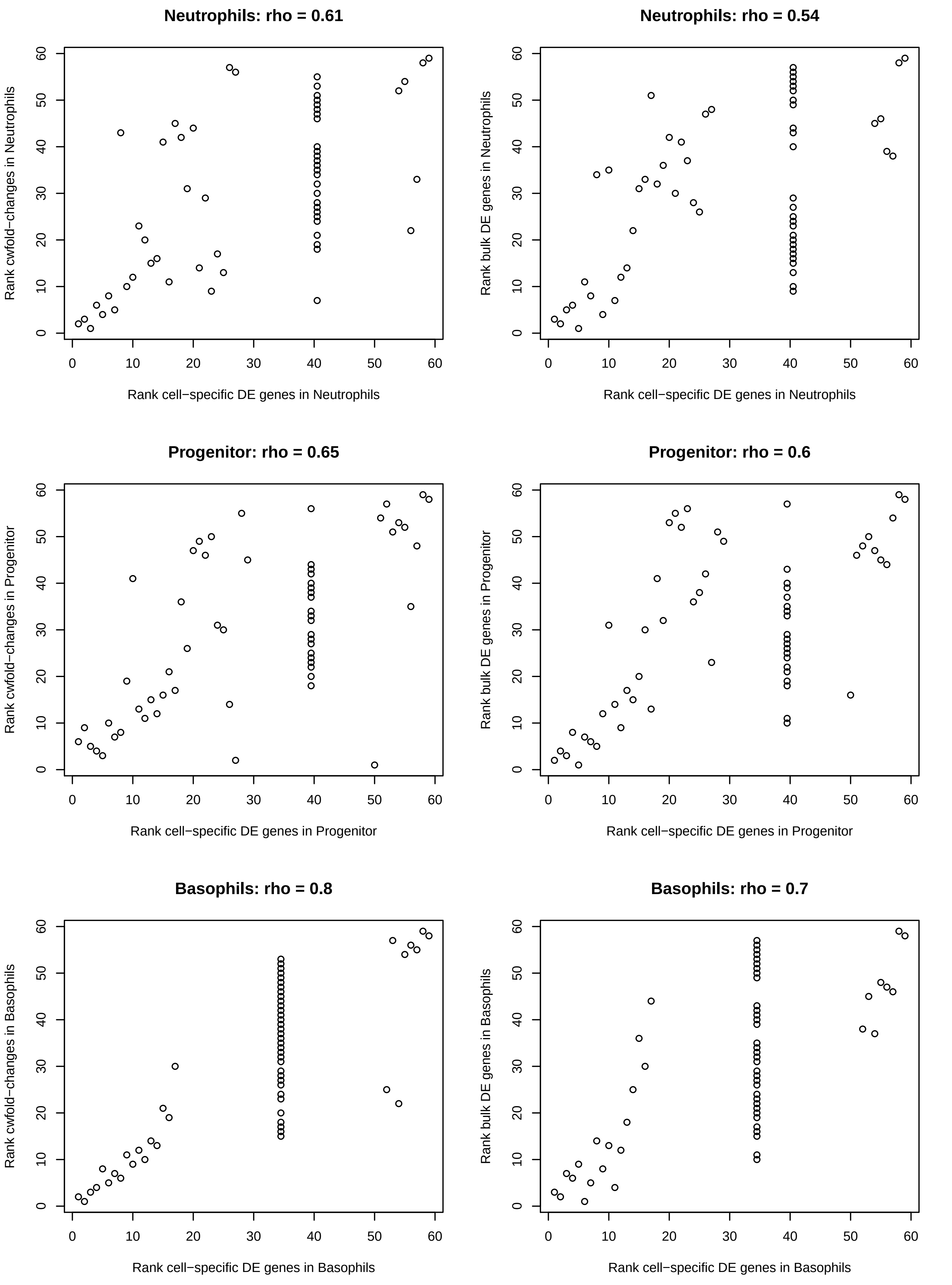
Neutrophils: rho = 0.61
Rank cwfold−changes in Neutrophils
Rank cell−specific DE genes in Neutrophils
Neutrophils: rho = 0.54
Rank bulk DE genes in Neutrophils
Rank cell−specific DE genes in Neutrophils
Progenitor: rho = 0.65
Rank cwfold−changes in Progenitor
Rank cell−specific DE genes in Progenitor
Progenitor: rho = 0.6
Rank bulk DE genes in Progenitor
Rank cell−specific DE genes in Progenitor
Basophils: rho = 0.8
Rank cwfold−changes in Basophils
Rank cell−specific DE genes in Basophils
Basophils: rho = 0.7
Rank bulk DE genes in Basophils
Rank cell−specific DE genes in Basophils

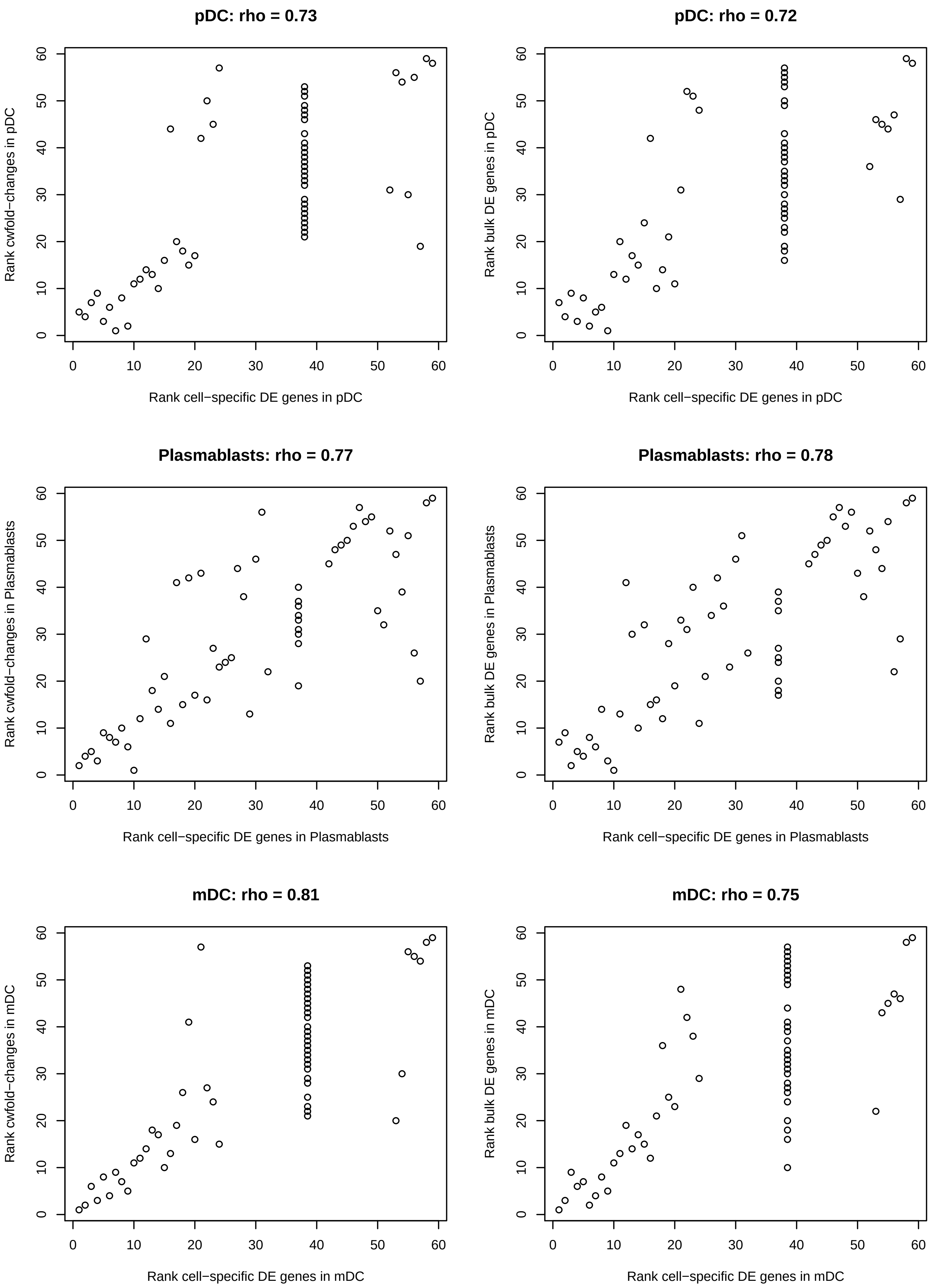
pDC: rho = 0.73
Rank cwfold−changes in pDC
Rank cell−specific DE genes in pDC
pDC: rho = 0.72
Rank bulk DE genes in pDC
Rank cell−specific DE genes in pDC
Plasmablasts: rho = 0.77
Rank cwfold−changes in Plasmablasts
Rank cell−specific DE genes in Plasmablasts
Plasmablasts: rho = 0.78
Rank bulk DE genes in Plasmablasts
Rank cell−specific DE genes in Plasmablasts
mDC: rho = 0.81
Rank cwfold−changes in mDC
Rank cell−specific DE genes in mDC
mDC: rho = 0.75
Rank bulk DE genes in mDC
Rank cell−specific DE genes in mDC

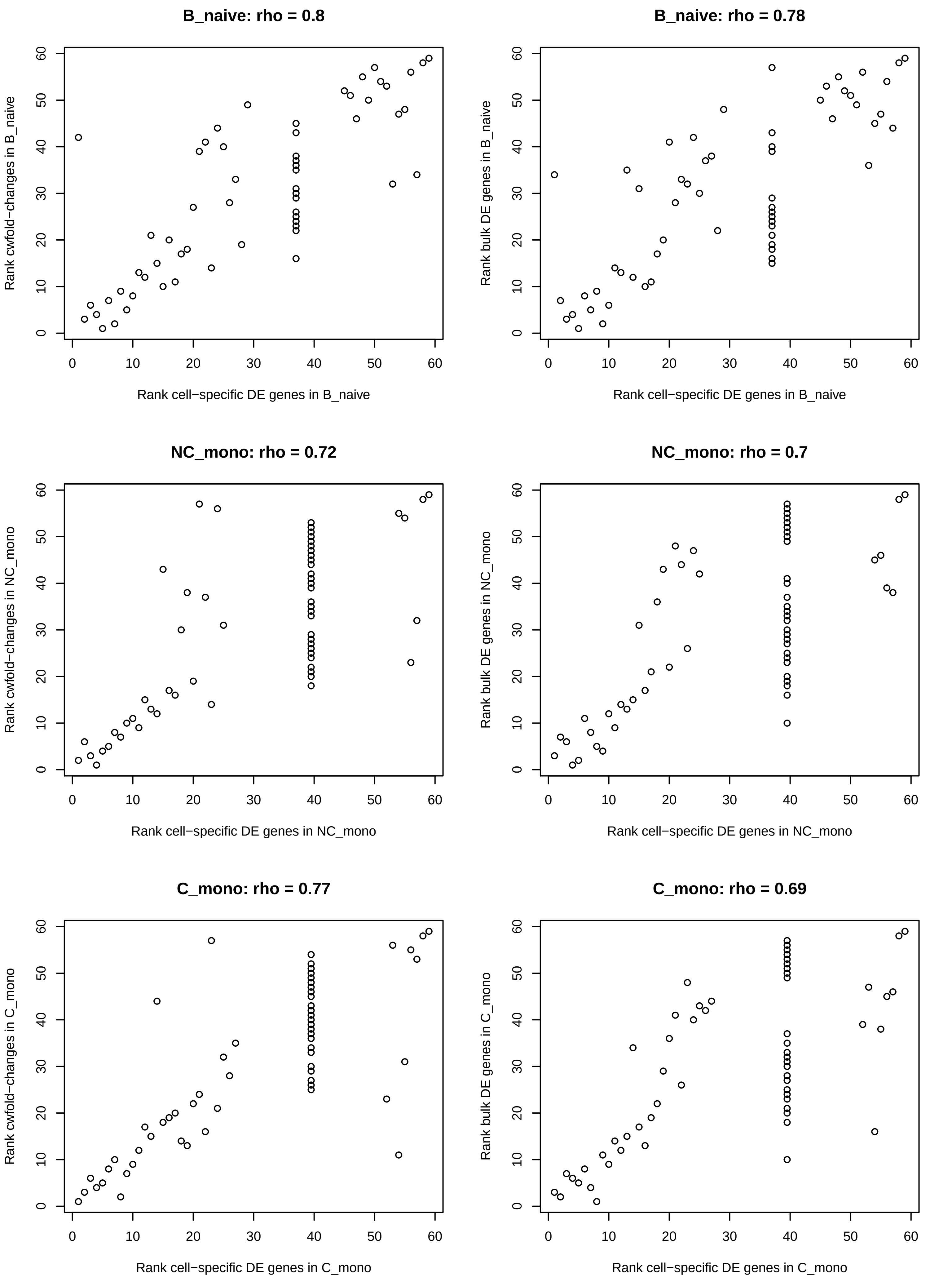
B_naive: rho = 0.8
Rank cwfold−changes in B_naive
Rank cell−specific DE genes in B_naive
B_naive: rho = 0.78
Rank bulk DE genes in B_naive
Rank cell−specific DE genes in B_naive
NC_mono: rho = 0.72
Rank cwfold−changes in NC_mono
Rank cell−specific DE genes in NC_mono
NC_mono: rho = 0.7
Rank bulk DE genes in NC_mono
Rank cell−specific DE genes in NC_mono
C_mono: rho = 0.77
Rank cwfold−changes in C_mono
Rank cell−specific DE genes in C_mono
C_mono: rho = 0.69
Rank bulk DE genes in C_mono
Rank cell−specific DE genes in C_mono

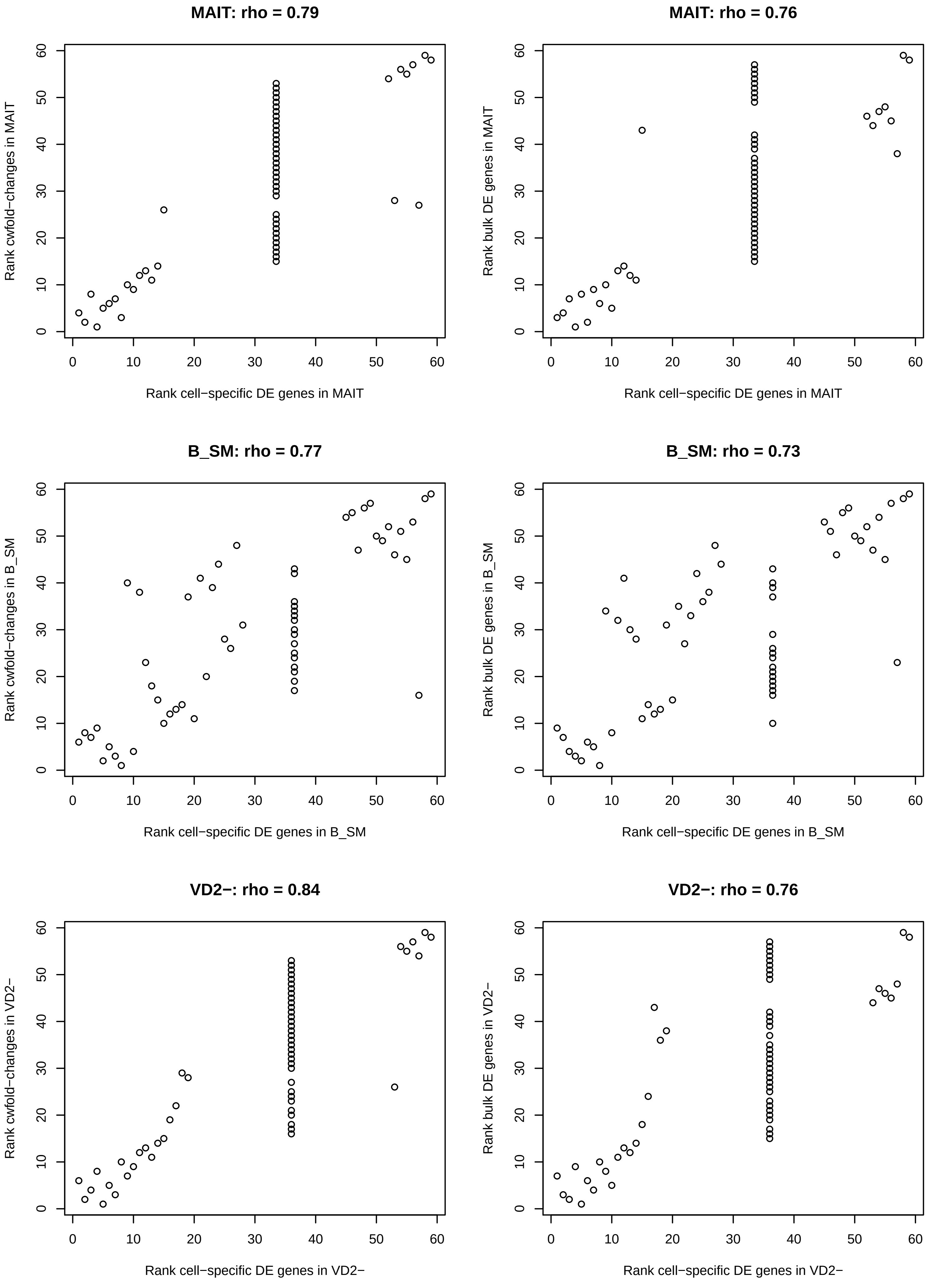
MAIT: rho = 0.79
Rank cwfold−changes in MAIT
Rank cell−specific DE genes in MAIT
MAIT: rho = 0.76
Rank bulk DE genes in MAIT
Rank cell−specific DE genes in MAIT
B_SM: rho = 0.77
Rank cwfold−changes in B_SM
Rank cell−specific DE genes in B_SM
B_SM: rho = 0.73
Rank bulk DE genes in B_SM
Rank cell−specific DE genes in B_SM
VD2−: rho = 0.84
Rank cwfold−changes in VD2−
Rank cell−specific DE genes in VD2−
VD2−: rho = 0.76
Rank bulk DE genes in VD2−
Rank cell−specific DE genes in VD2−

**Supplementary Figure 3. Scatterplots comparing increases in cell-type specificity after scMappR is applied.** The rank order comparing differentially expressed (DE) genes of sex differences within each cell-type (X-axis, all plots) to the rank order of cell-weighted fold changes (cwFold-changes) (Y-axis, left column) and DE genes in peripheral blood mononuclear cells (PBMC) (right, Y-axis, right column) are plotted. The stack of genes in the middle of each plot are genes that are not expressed in that cell-type. These genes are the same rank occurring directly in the middle of up- and down-regulated genes. Neutrophils = Neutrophils, Progenitor = Progenitor, Basophils = Basophils, pDC = Plasmacytoid dendritic cells, Plasmablast = Plasmablast, mDC = myeloid dendritic cells, B_naive = naive B cells, NC_mono = non-classical monocytes, C_mono = classical monocytes, MAIT = MAIT cells, B_SM = Switched memory B cells, VD2- = non-Vd2 gd T-cells.

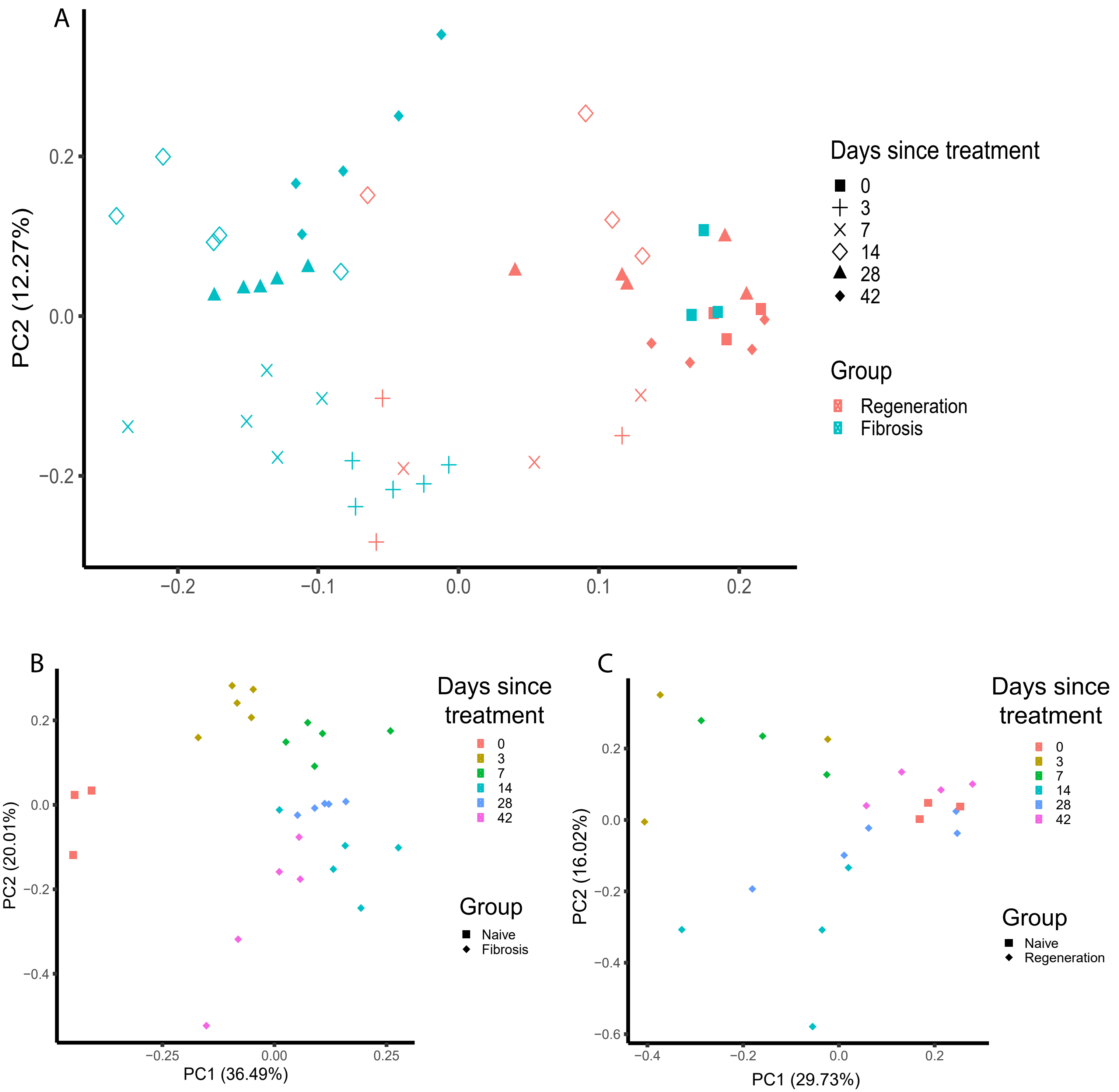


**Supplementary Figure 4. Principal component analysis (PCA) of bulk kidney RNA-seq samples in do Valle Duraes et al., 2020.** A) PCA of all samples, regeneration and fibrosis. Colour represents genotype (and also regeneration (all mutant) or fibrosis (all wildtype)). Shape represents time-point. B) PCA of the regeneration cohort, where shape represents naive or regeneration and colour represents time since injury. C) PCA of the fibrosis cohort, where shape represents naive (sham) or fibrosis and colour represents time since injury.

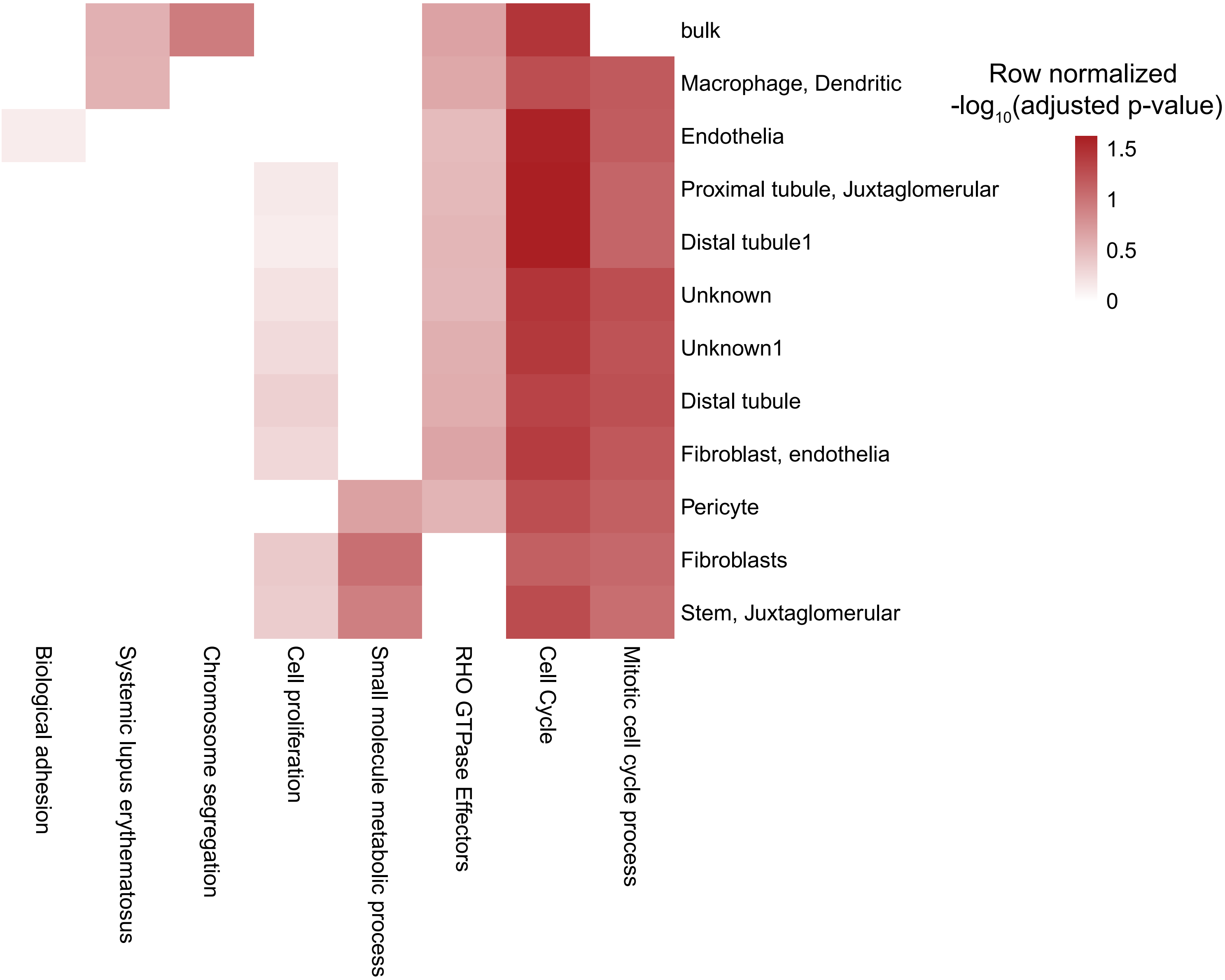


**Supplementary Figure 5. Cell-type normalized matrix of the top four most enriched pathways of differentially expressed genes re-ranked for each cell-type with scMappR.** Each row is a Gene Ontology, Reactome, or KEGG pathway identified with g:Profiler and is one of the top 4 pathways for at least one cell-type. Each column is a different cell-type. The heat of the heatmap is the -log10 of the FDR adjusted P-value for gene set enrichment. "Bulk" represents pathway enrichment before scMappR is completed.

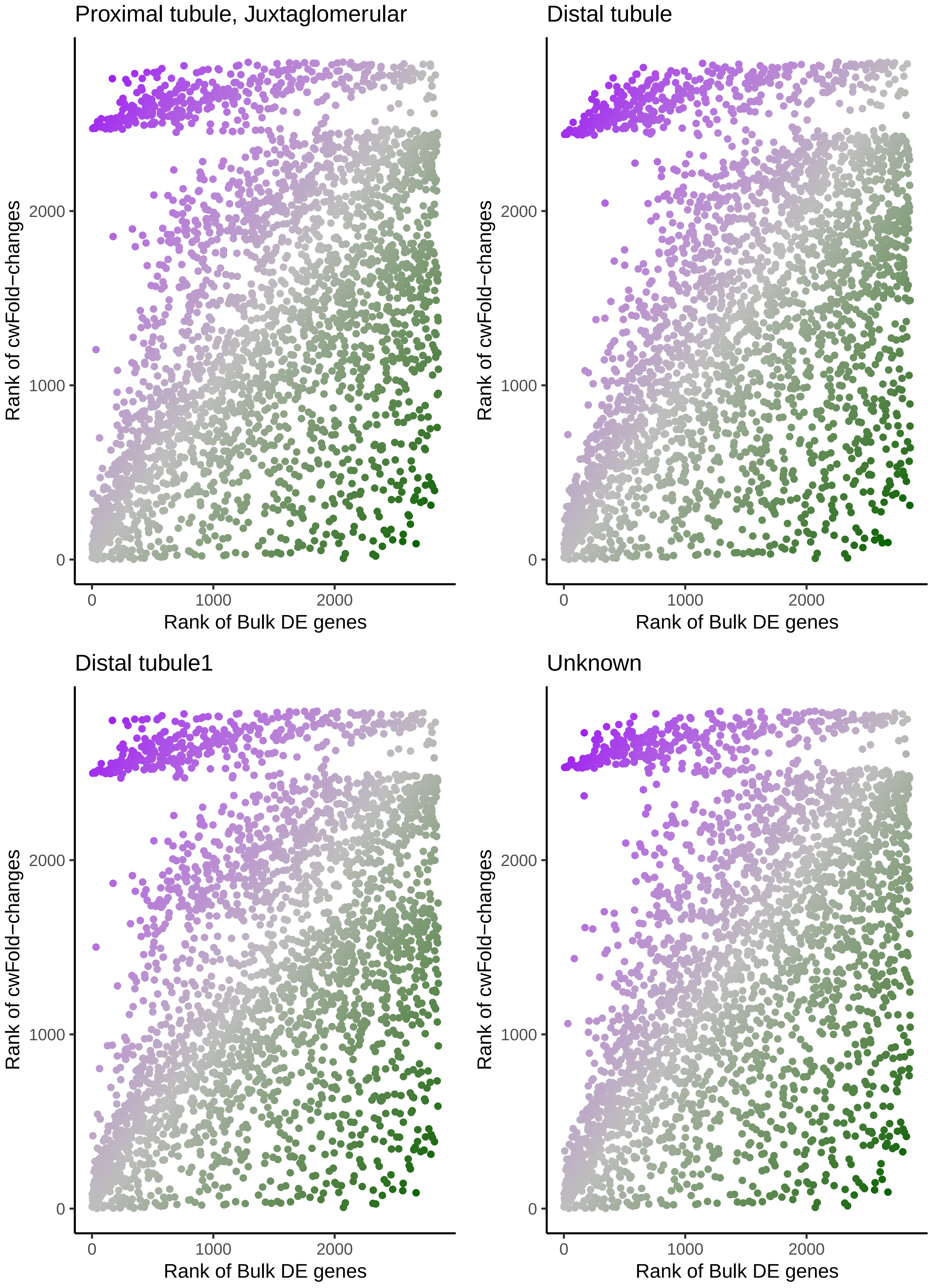
Proximal tubule, Juxtaglomerular
Distal tubule
Distal tubule1
Unknown
Rank of cwFold−changes
Rank of Bulk DE genes
0
1000
2000

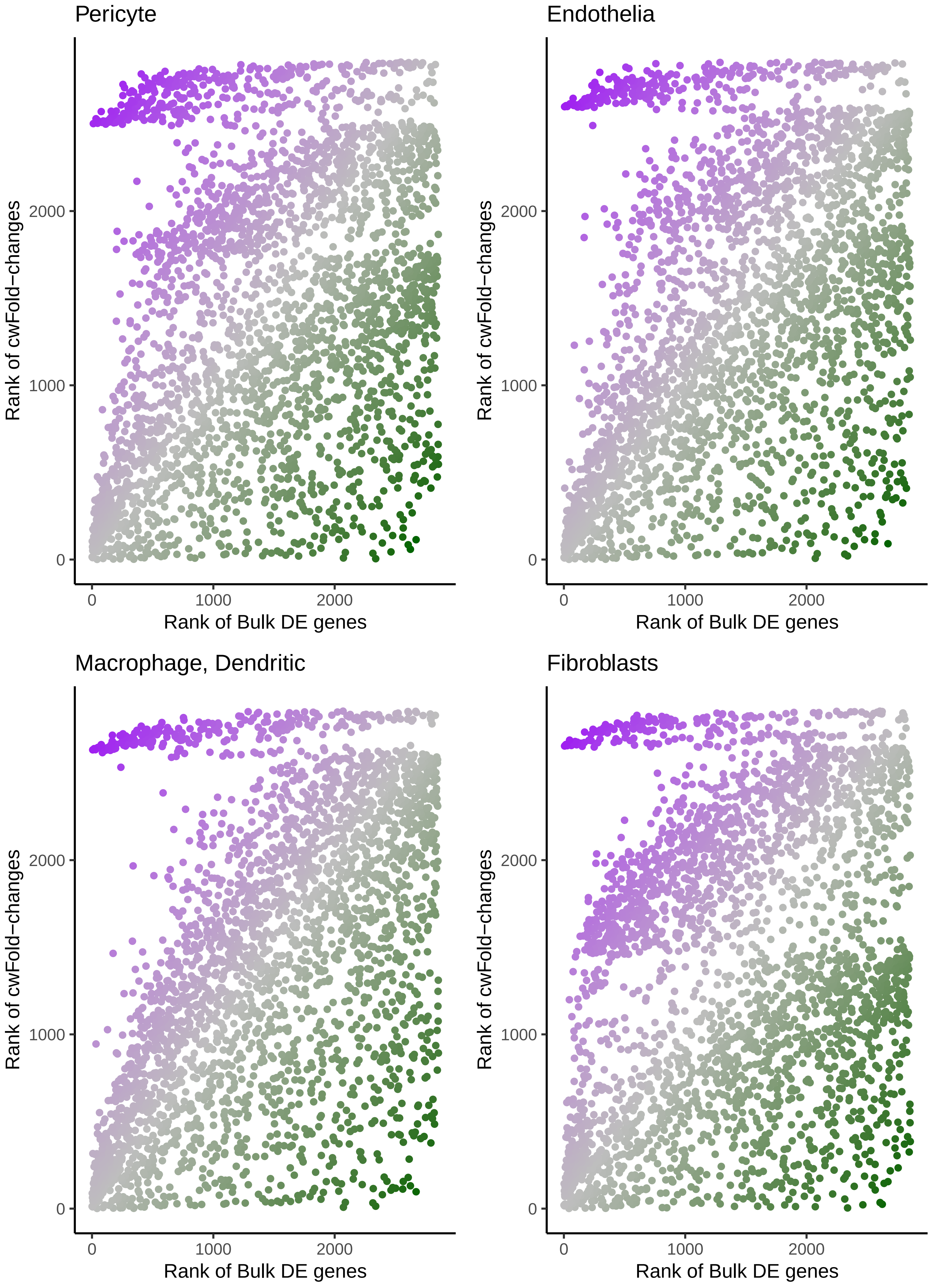

Pericyte
Endothelia
Macrophage, Dendritic
Fibroblasts
Rank of cwFold−changes
Rank of Bulk DE genes
0
1000
2000

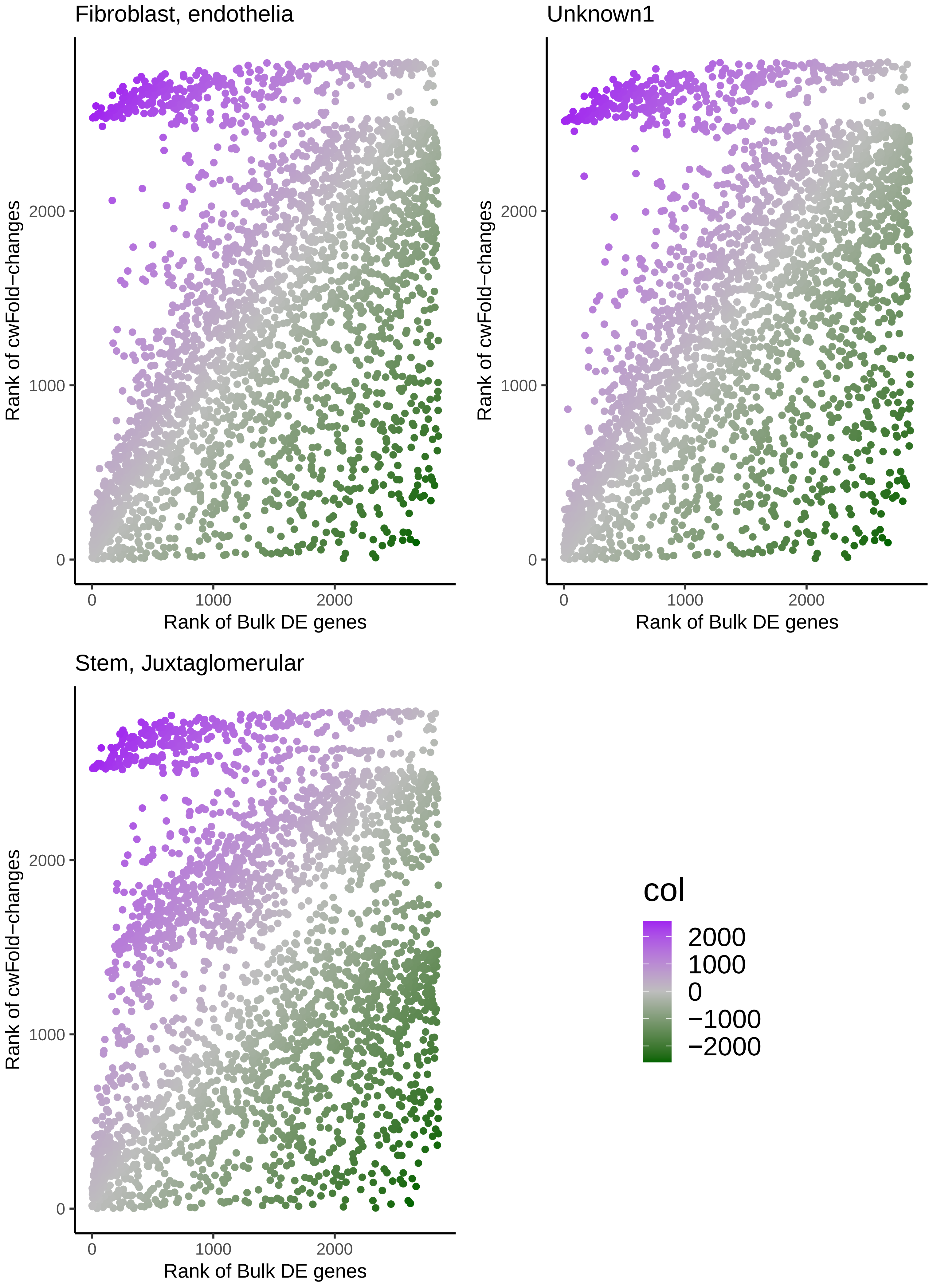

Fibroblast, endothelia
Unknown1
Stem, Juxtaglomerular
Rank of cwFold−changes
Rank of Bulk DE genes
0
1000
2000
col
2000
1000
0
−1000
−2000

**Supplementary Figure 6. Scatterplots comparing the ranks of differentially expressed genes between naive (day 0) and regeneration day 3 (day 3) before and after scMappR is applied.** The X-axis is the rank order of bulk differentially expressed genes and the Y-axis is the rank order of these genes after scMappR is applied. Genes in purplehave a higher rank in bulk and genes in green have a higher rank for that cell-type. The genes that have a low rank after scMappR is applied regardless of their initial rank (i.e. the relatively flat blue lines at the top) are the genes that are not expressed in that cell-type.

| Species | Studies | Total cell-types identified | Tissues | Signature matrices |
|---|---|---|---|---|
| Mouse | 122 | 4407 | 131 | 187 |
| Human | 46 | 1153 | 44 | 58 |

**Supplementary Table 1.** Summary of the signature matrices generated in scMappR at the time of publication.

| Cell-type | Number enriched pathways (adjusted p-value < 0.05) | Number of pathways not overlapping with bulk DEGs | Proportion of cwFold-change specific pathways | adjusted P-value for 'Immune Response' |
|---|---|---|---|---|
| Macrophage, dendritic | 137 | 77 | 0.562 | $1.62 \times 10^{-6}$ |
| Distal tubule1 | 126 | 67 | 0.532 | 0.00846 |
| Proximal tubule, juxtaglomerular | 127 | 71 | 0.559 | 0.0124 |
| Endothelia | 145 | 83 | 0.572 | 0.0259 |
| Distal tubule | 124 | 66 | 0.532 | 1.00 |
| Unknown | 125 | 67 | 0.536 | 1.00 |
| Pericyte | 127 | 71 | 0.559 | 1.00 |
| Fibroblasts | 133 | 78 | 0.586 | 1.00 |
| Fibroblast, endothelia | 128 | 68 | 0.531 | 1.00 |
| Unknown1 | 126 | 66 | 0.524 | 1.00 |
| Stem, juxtaglomerular | 133 | 77 | 0.579 | 1.00 |

**Supplementary Table 2. Number of enriched pathways of 2855 DEGs between naïve day 0 and regeneration day 3.** This table displays the overlap of the number of pathways from traditional bulk differential analysis, and from pathway analysis of the same DEGs re-ranked by cwFold-changes for each cell-type. It also displays the cell-type enrichment of the "Immune Response" gene ontology which was not enriched in the bulk pathway analysis.

| | Proximal tubule, juxtaglomerular | Endothelia | Macrophage, dendritic | Fibroblast, endothelia |
|---|---|---|---|---|
| Spearman's correlation of gene rank between 50 sample and 7 sample dataset | 1.00 | 1.00 | 1.00 | 1.00 |
| Ratio of average betewen 50 sample and 7 sample dataset | 1.00 | 1.00 | 1.00 | 1.00 |
| Average cell-type proportions in 50 sample dataset | 0.443 | 0.276 | 0.038 | 0.243 |
| Average cell-type proportions in 7 sample dataset | 0.431 | 0.289 | 0.029 | 0.251 |

**Supplementary Table 3. Comparison of the order of cwFold-changes, average cwFold-change value, and cell-type proportions between the 50 sample and 7 sample datasets.**

| SRA ID | Tissue | Label from CellMarker database | Label from Panglao database | Number of cell-type makers | Number of overlapping cell-type markers | Odds ratio | Adjusted p-value | Rank of enriched cell-types |
|---|---|---|---|---|---|---|---|---|
| SRA691388 | Breast epithelium | Neural progenitor cell | Gamma delta T cells | 141 | 107 | 3.55 | $3.28 \times 10^{-17}$ | 1 |
| SRA739096 | Lung | Neural progenitor cell | Gamma delta T cells | 141 | 107 | 3.55 | $3.26 \times 10^{-17}$ | 2 |
| SRA637291 | Left Ventricle | Neural progenitor cell | Gamma delta T cells | 171 | 113 | 3.09 | $3.09 \times 10^{-15}$ | 3 |
| SRA666404 | Cortical cells | Neural progenitor cell | Gamma delta T cells | 171 | 113 | 3.09 | $2.22 \times 10^{-15}$ | 4 |
| SRA728071 | Foam cells from aortas | Neural progenitor cell | Gamma delta T cells | 141 | 101 | 3.34 | $2.22 \times 10^{-15}$ | 5 |
| SRA797009 | Spleen | Neural progenitor cell | Gamma delta T cells | 141 | 101 | 3.34 | $2.22 \times 10^{-15}$ | 6 |
| SRA665153 | Epithelial cells in airway luminal surface | Neural progenitor cell | Gamma delta T cells | 99 | 79 | 3.71 | $1.25 \times 10^{-13}$ | 7 |
| SRA739096 | Lung | Neural progenitor cell | Gamma delta T cells | 170 | 107 | 2.94 | $1.25 \times 10^{-13}$ | 8 |
| SRA729910 | Lung airway epithelial cells | Neural progenitor cell | Gamma delta T cells | 216 | 123 | 2.66 | $1.75 \times 10^{-13}$ | 9 |
| SRA796267 | SVZ-derived neural stem cells | Neural progenitor cell | Gamma delta T cells | 216 | 123 | 2.66 | $1.75 \times 10^{-13}$ | 10 |
| SRA772115 | Kidney | Cycling basal cell | Gamma delta T cells | 79 | 63 | 3.69 | $4.37 \times 10^{-11}$ | 22 |

**Supplementary Table 4. Over-representation of cell-type markers of consistently processed scRNA-seq data in over 100 mouse tissues when inputting 2855 DEGs between naïve (day 0) and regeneration (day 3) bulk kidneys.**

| Cell Type | Total number of cell-type markers in *Tabula Muris*, 2018 | Upregulated DEGs | | | Downregulated DEGs | | |
|---|---|---|---|---|---|---|---|
| | | Odds ratio | Adjusted p-value | Number of genes | Odds ratio | Adjusted p-value | Number of genes |
| Macrophage, dendritic | 430 | 1.88 | $1.43 \times 10^{-5}$* | 151 | 0.333 | $4.23 \times 10^{-5}$* | 18 |
| Fibroblasts | 548 | 1.53 | 0.00384* | 166 | 0.442 | $4.10 \times 10^{-4}$* | 31 |
| Distal tubule | 48 | 0.252 | 0.0309* | 3 | 2.21 | 0.0901 | 10 |
| Endothelia | 560 | 0.720 | 0.0309* | 110 | 0.672 | 0.0901 | 43 |
| Stem, juxtaglomerular | 23 | 1.83 | 0.264 | 10 | 0.437 | 1.00 | 1 |
| Distal tubule1 | 111 | 0.691 | 0.305 | 19 | 1.98 | 0.0545 | 20 |
| Proximal tubule, juxtaglomerular | 17 | 0.242 | 0.356 | 1 | 1.81 | 0.761 | 3 |
| Pericyte | 49 | 1.37 | 0.367 | 16 | 0.823 | 1.00 | 4 |
| Unknown1 | 3 | 1.38 | 0.710 | 1 | 0.00 | 1.00 | 0 |
| Fibroblasts, endothelia | 43 | 1.06 | 0.948 | 11 | 0.943 | 1.00 | 4 |
| Unknown | 49 | 1.03 | 1.00 | 12 | | 1.00 | 4 |

**Supplementary Table 5. Over- and under-representation of kidney cell-type markers from scRNA-seq data generated by *Tabula Muris* when inputting 2855 DEGs between naïve (day 0) and regeneration (day 3) bulk kidneys.**
